# The contributions of DNA accessibility and transcription factor occupancy to enhancer activity during cellular differentiation

**DOI:** 10.1101/2023.02.22.529579

**Authors:** Trevor Long, Tapas Bhattacharyya, Andrea Repele, Madison Naylor, Sunil Nooti, Shawn Krueger, Manu

## Abstract

The upregulation of gene expression by enhancers depends upon the interplay between the binding of sequence-specific transcription factors (TFs) and DNA accessibility. DNA accessibility is thought to limit the ability of TFs to bind to their sites, while TFs can increase accessibility to recruit additional factors that upregulate gene expression. Given this interplay, the causative regulatory events underlying the modulation of gene expression during cellular differentiation remain unknown for the vast majority of genes. We investigated the binding-site resolution dynamics of DNA accessibility and the expression dynamics of the enhancers of an important neutrophil gene, *Cebpa*, during macrophage-neutrophil differentiation. Reporter genes were integrated in a site-specific manner in PUER cells, which are progenitors that can be differentiated into neutrophils or macrophages *in vitro* by activating the pan-leukocyte TF PU.1. Time series data show that two enhancers upregulate reporter expression during the first 48 hours of neutrophil differentiation. Surprisingly, there is little or no increase in the total accessibility, measured by ATAC-Seq, of the enhancers during the same time period. Conversely, total accessibility peaks 96 hrs after PU.1 activation—consistent with its role as a pioneer—but the enhancers do not upregulate gene expression. Combining deeply sequenced ATAC-Seq data with a new bias-correction method allowed the profiling of accessibility at single-nucleotide resolution and revealed protected regions in the enhancers that match all previously characterized TF binding sites and ChIP-Seq data. Although the accessibility of most positions does not change during early differentiation, that of positions neighboring TF binding sites, an indicator of TF occupancy, did increase significantly. The localized accessibility changes are limited to nucleotides neighboring C/EBP-family TF binding sites, showing that the upregulation of enhancer activity during early differentiation is driven by C/EBP-family TF binding. These results show that increasing the total accessibility of enhancers is not sufficient for upregulating their activity and other events such as TF binding are necessary for upregulation. Also, TF binding can cause upregulation without a perceptible increase in total accessibility. Finally, this study demonstrates the feasibility of comprehensively mapping individual TF binding sites as footprints using high coverage ATAC-Seq and inferring the sequence of events in gene regulation by combining with time-series gene expression data.

## 1 Introduction

Metazoan gene expression is regulated intricately in time during cellular differentiation [1–3]. The cell-type specific temporal pattern of a gene’s expression is encoded in DNA as clusters of transcription factor (TF) binding sites called *cis*-regulatory modules (CRMs) or enhancers [1, 4]. TFs bound to enhancers recruit co-activators or co-repressors that interact with the RNA polymerase holoenzyme complex directly or through the Mediator complex [5, 6]. In addition to the action of sequence-specific TFs, the accessibility of enhancer DNA, which is inversely related to nucleosome occupancy, has been linked to enhancers’ ability to drive higher levels of gene expression [7–12]. While enhancer accessibility is thought to influence the occupancy of sequence-specific TFs, bound TFs also regulate accessibility by either actively recruiting chromatin remodeling enzymes, or passively by competing with nucleosomes [9, 13–15]. Furthermore, enhancers, especially those regulating developmental genes, have complex regulation and some are regulated by 6–8 TFs binding to multiple sites for each TF [16–21]. Consequently, the causative regulatory events underlying the modulation of gene expression in time remain unknown for the vast majority of genes.

*CCAAT/Enhancer binding protein, α* (*Cebpa*) encodes a TF that is necessary for neutrophil development [22] as well as the specification of hepatocytes and adipocytes [23, 24]. During hematopoiesis, *Cebpa* is expressed in hematopoietic stem cells, granulocyte-monocyte progenitors (GMPs), neutrophils, and macrophages (`http: //biogps.org/gene/12606`; [25, 26]). Although the most apparent hematopoietic phenotype of *Cebpa*^−^*^/^*^−^ mice is neutropenia [22], *Cebpa* also has a role in specifying macrophages. *Cebpa* is expressed at intermediate and high levels in macrophages and neutrophils respectively and the cell-fate decision is thought to depend on the ratio of PU.1, a TF necessary for all white-blood cell lineages [27], and C/EBP*α* expression levels [28].

Reflecting it’s functions in multiple tissues, *Cebpa* has a complex gene regulatory architecture. The promoter is bound by C/EBP family, Usf1, NF-Y family, Myc, ZNF143, and two other unknown TFs [23, 29]. In addition to the promoter, *Cebpa* is regulated by three enhancers located 8kb, 31kb, and 37kb downstream of the *Cebpa* transcription start site (TSS) in the mouse genome (Fig. 1A). The enhancer located at 37kb is bound by PU.1, other Ets TFs, C/EBP, Runx1, SCL, Gata2, and Myb [17, 18, 30, 31] in myeloid cells. The enhancer at 8kb is bound by C/EBP and Gfi1 [18, 31, 32] and the third enhancer is bound by PU.1 [18, 31]. While the binding sites and identities of the TFs regulating *Cebpa* enhancers have been established, it is not understood how the regulatory contributions of these TFs modulate *Cebpa*’s gene expression during myeloid differentiation. In particular, neither the dynamics of enhancer-driven expression, nor the dynamics of enhancer chromatin state have been characterized.

**Figure 1:**
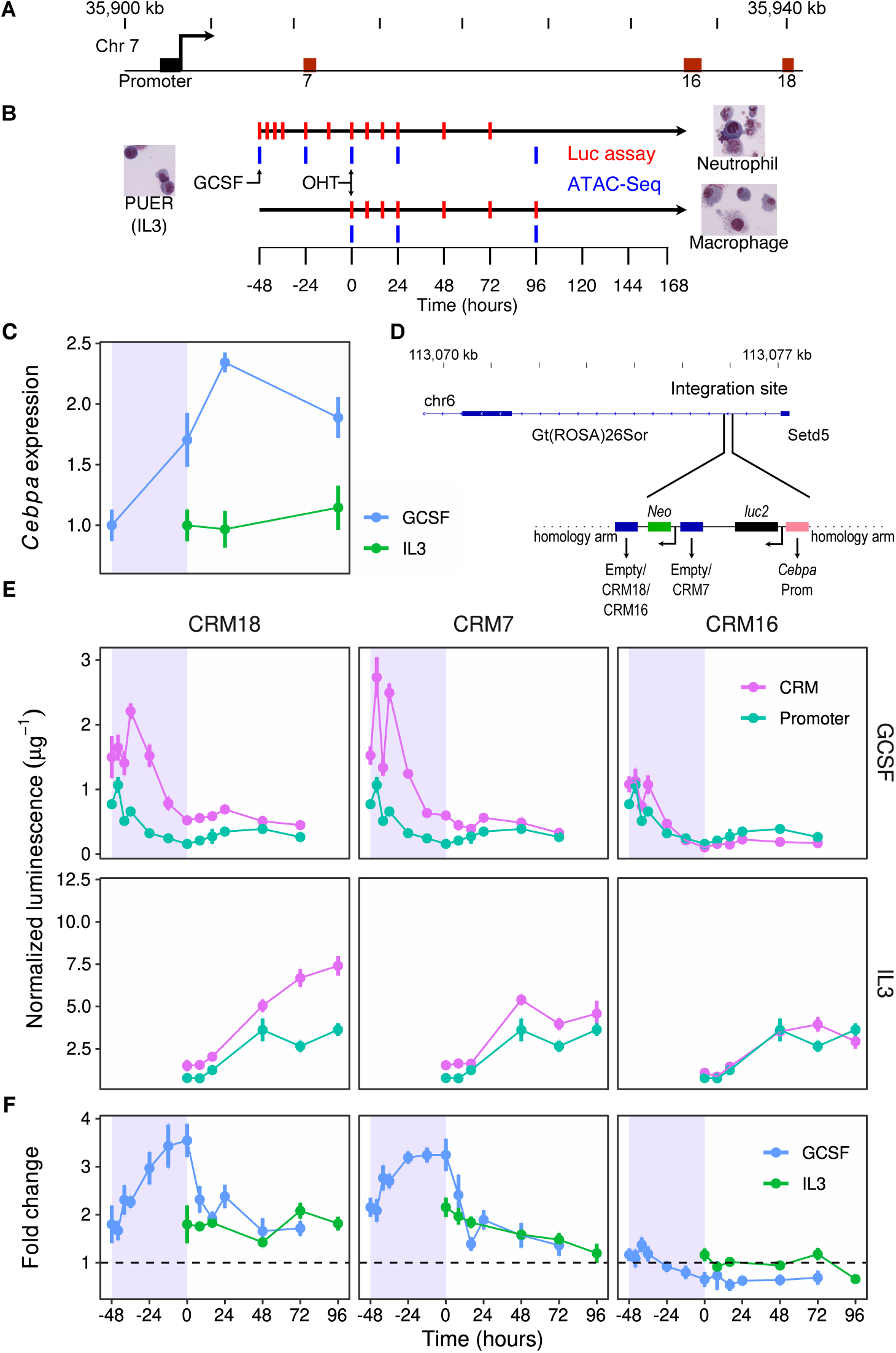
Expression driven by *Cebpa* CRMs during PUER cell differentiation. **A**. A 40kb region of chromosome 7 showing the *Cebpa* transcription start site (TSS), promoter, and CRMs (enhancers) 7, 16, and 18. **B**. PUER cells may be differentiated into macrophages over a 7-day period by inducing with OHT in IL3 medium. Substituting GCSF for IL3 48 hours prior to OHT induction results in neutrophil differentiation. The red and blue tick marks indicate timepoints at which Luciferase assays and ATAC-Seq were conducted respectively. **C**. The time course of *Cebpa* expression relative to *hprt* measured by RT-qPCR during PUER differentiation. Blue shaded region shows period of GCSF pre-treatment prior to induction with OHT during neutrophil differentiation. **D**. Site-specific knock in of Luciferase reporters into intron 1 of the ROSA26 locus by CRISPR/Cas9 HDR. *luc2* gene expression was driven by the *Cebpa* promoter (orange) alone or along with CRM 7, 16, or 18. **E**. Time series of Luciferase expression normalized to total protein measured in *µ*g of BSA. Expression in GCSF and IL3 conditions is shown in the top and bottom panels respectively. The expression driven by the promoter alone is shown in green, while the expression of reporters bearing CRM 18, 7, or 16 along with the promoter are shown in magenta in the left, middle, or right panels respectively. **F**. Fold change in the expression of enhancer-bearing reporters relative to the promoter at each time point.

We profiled DNA accessibility at binding-site resolution as well as the dynamics of both accessibility and gene expression of the *Cebpa* enhancers during the differentiation of an *in vitro* model of myeloid development. PU.1^−^*^/^*^−^ cells from mouse bone marrow carrying an *Spi1* (PU.1) transgene fused to the ligand-binding domain of the estrogen receptor [PUER; 33] can be inducibly differentiated into macrophages or neutrophils in IL3 or GCSF respectively [Fig. 1B; 18, 28, 31, 34]. *Cebpa* gene expression is upregulated by GCSF treatment during neutrophil differentiation but not during macrophage differentiation [Fig. 1C; 18, 28]. In order to measure the upregulation driven by each enhancer at a high temporal resolution, we used CRISPR/Cas9 to knock in Luciferase reporters containing the *Cebpa* promoter either alone or together with one of the enhancers, CRM 7 (+8kb), 16 (+31kb), or 18 (+37kb), in the same site in the ROSA26 locus in a biallelic manner (Fig. 1D). We measured the activity of each enhancer as the fold change in gene expression relative to the promoter, both in bulk and at the single-cell level using a flow cytometry-based method we developed for the purpose.

The temporal pattern of the fold change of CRMs 7 and 18 matches that of the endogenous gene, showing that they are primarily responsible for the GCSF-dependent upregulation during neutrophil differentiation. Pooling ATAC-Seq samples allowed us to achieve an average depth of coverage of ∼50 Tn5 cuts per nucleotide in the accessible regions of the genome, furnishing a very high resolution picture of intra-enhancer accessibility. These high resolution profiles contained footprints that match all previously characterized binding sites and overlap ChIP signal peaks of the cognate TFs. While enhancer activity and total accessibility are correlated, being higher in GCSF conditions than in IL3, there isn’t a consistent causal connection between the two. *Cebpa* expression and the fold change of CRMs 7 and 18 peak early during the differentiation and change relatively little at later time points. The total accessibility of the enhancers changes little during early differentiation and peaks at later timepoints after PU.1 induction, which is consistent with PU.1’s role as a pioneer factor [35–38]. The high resolution of the DNA accessibility profiles allowed us to examine the accessibility of individual nucleotides adjacent to TF binding sites, which is an indicator of TF occupancy [39, 40]. The accessibility of nucleotides neighboring specific C/EBP-family TF binding sites increases at early timepoints, showing that enhancer upregulation is driven by increased TF occupancy. These results support a model in which *Cebpa* is upregulated by GCSF treatment due to increased binding of sequence-specific TFs, and the later increase in total accessibility of the enhancers, while being a consequence of PU.1 binding, does not result in increased enhancer activity.

## 2 Results

### 2.1 Enhancer Activity

We utilized CRISPR/Cas9 homology directed repair (HDR) with a “double-cut” donor strategy [41] to create transgenic PUER cell lines bearing synthetic reporter loci in the ROSA26 locus (Fig. 1D). We created a line with the *Cebpa* promoter driving *luc2* reporter gene transcription but lacking an enhancer (promoter line), and lines bearing an enhancer, CRM 7, 16, or 18, in addition to the promoter (referred to as CRM 7, 16, and 18 lines respectively). Our experimental strategy (Section 4) ensured that the synthetic loci were integrated into the same location in a biallelic manner and that the integration was seamless. Multiple biallelic clones derived from independent HDR events had reporter expression within 30% of each other (Fig. S1) and a representative clone was selected for further time series experiments (Section 4).

We measured reporter expression during the differentiation of PUER cells by Luciferase assay at several time points. PUER cells are maintained in IL3 medium and can be differentiated into macrophage-like cells over a period of 7 days by activating the PU.1-estrogen receptor fusion protein with 4-hydroxy-tamoxifen (OHT) [18, 28, 31]. Replacing IL3 medium with GCSF medium for 48 hours prior to OHT induction causes PUER cells to differentiate into neutrophils over a 7-day period. We regard OHT induction as the initiation of differentiation so that undifferentiated PUER cells correspond to the 0h time point in IL3 conditions and the -48h time point GCSF conditions (Fig. 1B).

The mRNA and protein expression of the endogenous *Cebpa* gene has a characteristic temporal profile, being upregulated about two-fold relative to undifferentiated cells during the 48-hour pre-treatment with GCSF, peaking at 24 hours after OHT induction and then declining subsequently [Fig. 1C; 18, 28]. In contrast, *Cebpa* expression is relatively constant in IL3 conditions so that the GCSF response accounts for the neutrophil-specific upregulation of the gene.

During neutrophil differentiation, reporter gene expression per cell (Fig. 1E), estimated by normalizing the observed Luciferase luminescence to total protein, shows a transient upregulation peaking during the first 12 hours of GCSF pre-treatment for all the reporter genes. CRMs 18 and 7 have the largest effect, peaking 50% and 80% over their levels in undifferentiated cells respectively, while the promoter peaks at a 40% increase over its level in undifferentiated cells. The activity of the CRMs may be characterized by computing the fold change in expression relative to the promoter. As expected, all three enhancers drove reporter expression at levels higher than the promoter in undifferentiated PUER cells [Fig. 1F; 18], with CRMs 7 and 18 having a fold change of ∼2. The fold change of CRMs 7 and 18 increases to ∼3.5 during the GCSF pre-treatment and declines upon OHT induction, staying above 1 at all times. In contrast, the fold change of CRM 16 declines to less than 1 during neutrophil differentiation, suggesting a dominant negative effect on the transcription of the promoter. We checked these measurements by an independent method we developed to assay reporter gene expression at the single-cell level using flow cytometry (Section 4). The single-cell reporter data have unimodal distributions of Luciferase expression (Fig. S2B) showing that gene expression is not heterogeneous. The time series of median expression (Fig. S2C) agree with the bulk measurements. Overall, the temporal pattern of the fold change of CRMs 7 and 18 (Fig. S2D) matches that of endogenous *Cebpa* during GCSF pre-treatment, showing that these two CRMs are primarily responsible for the gene’s neutrophil-specific upregulation.

OHT induction in IL3 conditions causes a comparable upregulation of all the reporter genes, so that the fold change remains relatively constant for CRMs 16 and 18 and declines from ∼2 to ∼1 for CRM 7 (Fig. 1E). As in GCSF conditions, the temporal pattern of fold change matches the relatively constant expression of the endogenous gene in IL3 conditions.

### 2.2 Accessibility Landscape of *Cebpa*

In order to discover the regulatory mechanisms driving the temporal patterns of reporter expression, we profiled the accessibility of the *Cebpa* locus with deep sequencing at several time points during the differentiation of PUER cells into macrophages and neutrophils. The accessibility of regulatory regions is regarded as a determinant of gene expression since it could influence the occupancy of TF binding sites [42, 43]. Conversely, TFs influence the accessibility of enhancers by competing with and displacing nucleosomes [15, 44, 45]. Profiling accessibility with deep sequencing has the potential to reveal TF binding sites as footprints [12, 40, 46–49] and the *cis*-regulatory logic of *Cebpa* regulatory regions.

We sampled cells in duplicate at multiple stages of PUER differentiation (Fig. 1B) and profiled their DNA accessibility using a variation of the classical ATAC-seq method [12] known as Fast-ATAC [50]. Following sampling, DNA libraries were prepared following the Fast-ATAC protocol (Section 4) and were sequenced to approximately 300 million reads/sample with paired-end sequencing. The sequences were then filtered, trimmed, and aligned to the *mm10* genome. We counted each read as one Tn5 cut. Tn5 leaves a 9bp 5’ overhang [Fig. S3; 51], so that the 5’ ends of reads resulting from the same transposition event are mapped to different locations. We shifted the position of the cut to the center of the overhang by adding +4bp/-4bp if the read mapped to +/-strand respectively (Fig. S3B,C). Tn5 transposition is not uniformly random on naked DNA and these biases can obscure TF footprints. Methods for correcting bias either average over 50–100bp [52] and thus do not correct bias at single-nucleotide resolution or do not average and amplify noise at low accessibility sites [39]. We developed a new method that overcomes these limitations (Section 4 and Fig. S4) in estimating bias from purified mouse genomic DNA ATAC-Seq libraries [53–56] and correcting it. Finally, corrected Tn5 cut counts were normalized to each sample’s library size.

The accessibility profile of a 110kb region centered on the *Cebpa* TSS (*Cebpa locus*) is highly correlated between replicates (Fig. 2B) and time points (Fig. 2A). Neither newly accessible regions appear nor are any accessible regions lost during differentiation. Instead, we observed relatively smooth quantitative changes in the accessibility of different regions in the locus (Fig. 2A), which is consistent with similar analyses in other cell types [20]. In undifferentiated PUER cells, the *Cebpa* promoter was the most accessible region, followed in descending order by CRM 18, 7, and 16. Interestingly, while *Cebpa* expression in GCSF conditions is roughly two-fold its value in IL3 (Fig. 1C), the accessibility of the *Cebpa* promoter is comparable between the conditions. The accessibility of two enhancers, CRMs 7 and 18, both increased approximately 2-fold during differentiation in neutrophil conditions suggesting correlation between the upregulation of *Cebpa* expression and the chromatin state of the enhancers.

**Figure 2:**
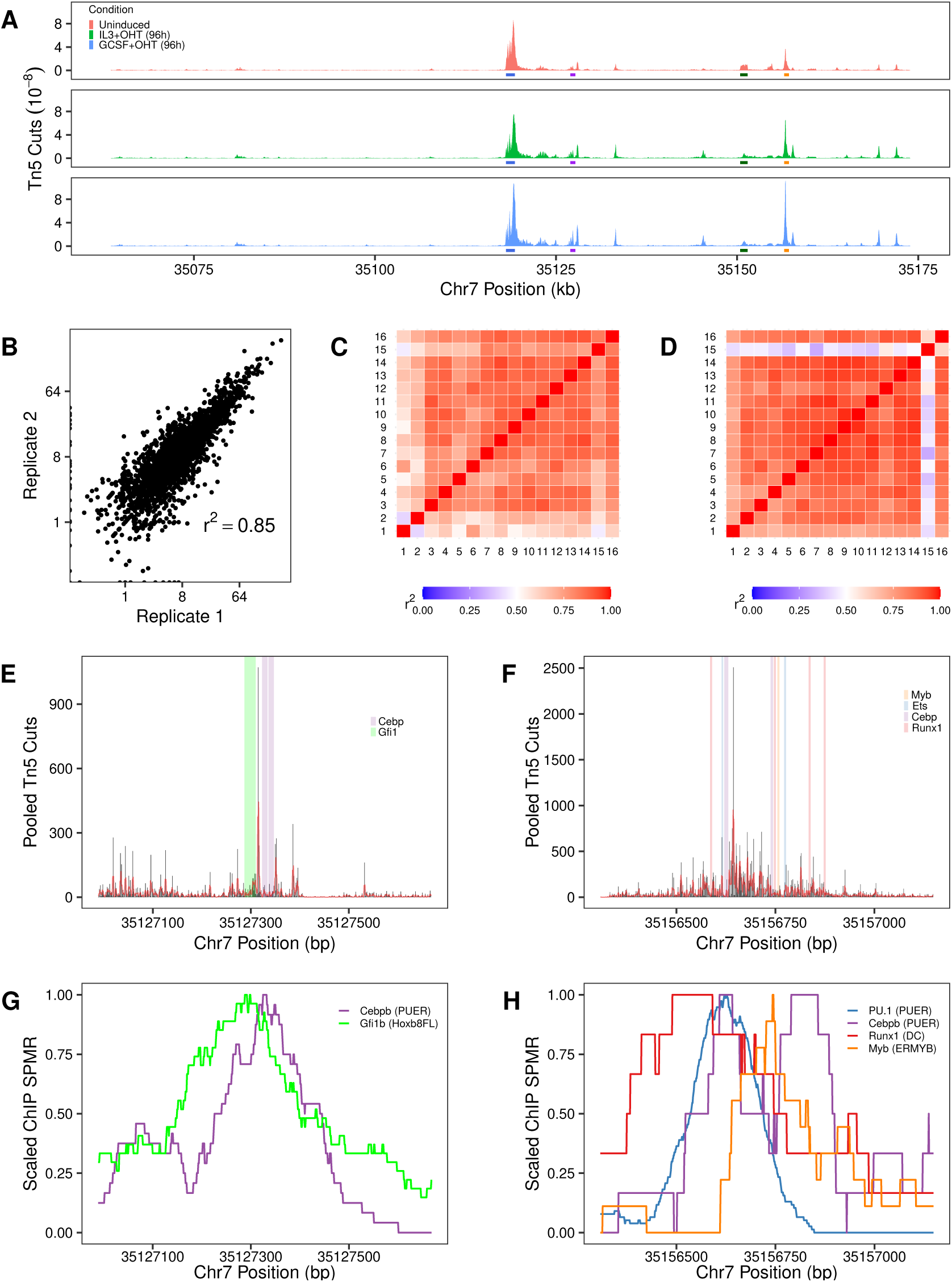
Locus-and enhancer-level accessibility profiles. **A**. The chromatin accessibility profile of a 110kb region (locus) centered on the *Cebpa* TSS in uninduced PUER cells (red), 96 hrs after OHT induction in IL3 (green, macrophage), and 96 hrs after OHT induction in GCSF (blue, neutrophil). **B**. Tn5 cuts at each nucleotide in the 110kb locus averaged with a 100bp sliding window. The two IL3+OHT (96h) replicates are compared. **C**. Pearson correlation coefficients of Tn5 counts in CRM 7, averaged with a 10bp sliding window, computed between each pair of samples. **D**. Pearson correlation coefficients of Tn5 counts in CRM 18, averaged with a 10bp sliding window, computed between each pair of samples. **E**. Pooled single-nucleotide accessibility profile of CRM 7. Black bars are the sum of Tn5 counts over all the samples. Red line is a 3bp sliding-window average. Shaded regions show empirically verified known binding sites. **F**. Pooled single-nucleotide accessibility profile of CRM 18. **G**. Scaled ChIP-Seq signal of C/EBP*β* in PUER cells (GSM538010) and Gfi1b in Hoxb8FL cells (GSM2231904) in CRM 7. **H**. Scaled ChIP-seq signal of C/EBP*β* (GSM538010) and PU.1 (GSM538004) in PUER cells, Runx1 in dendritic cells (GSM881115), and Myb in ERMYB cells (GSM549341) in CRM 18.

### 2.3 *Cebpa* Enhancer Structure

To further understand the regulatory logic of *Cebpa*’s enhancers, we analyzed their accessibility profiles at high resolution. The accessibility profiles of the promoter, CRM 7, and CRM 18 were highly correlated between samples (Fig. 2C,D and S5A), similar to the 110kb region. CRM 16 is an exception, having low correlation between samples (Fig. S5B), which could be the result of the dynamic changes in profile during differentiation (Fig. S14) or noise due to relatively lower accessibility (see below). The high correlation between samples suggests that the chromatin structure of the promoter and CRMs 7 and 18 remains relatively constant and that the enhancers are bound at the same sites throughout differentiation. The high correlation of CRMs 7 and 18 and the promoter allowed us to pool the samples by summing the Tn5 cuts at each nucleotide, resulting in a very high depth of coverage in the accessible regions of the genome. For example, after pooling, CRM 18 has a total of 39,691 reads over its 642bp length, resulting in a coverage of ∼62 reads/nucleotide. The pooled accessibility profile of the enhancers (Fig. 2E,F and S5C,D) consists of many short high-accessibility “islands” 1–5bp in length interspersed between 5–50bp-long regions having nearly zero accessibility. Such high-low-high accessibility patterns are regarded as TF footprints [12, 52], although there has been mixed success in detecting them at an individual level [39, 40, 52].

To determine whether these high-low-high patterns are, in fact, TF binding sites, we mapped previously characterized high-confidence binding sites to each enhancer. These sites have been identified and validated previously with multiple methods such as DNase I footprinting, EMSA, ChIP, and site-directed mutagenesis coupled to reporter assays [17, 18, 23, 31, 57]. All previously identified binding sites overlapped with the footprints (Fig. 2E,F and Fig. S5C,D). CRM 7 has protected binding sites for both Gfi1 and C/EBP-family TFs while CRM 18 has protection by a multitude of TFs including Myb, Ets-family, C/EBP-family, and Runx1, showing that these sites are occupied in PUER cells. We further validated the footprints against publicly available ChIP-Seq datasets, where available, for the TFs predicted to bind them, PU.1 and C/EBP*β* [PUER; 58], Runx1 [dendritic cells; 59], Myb [ERMYB; 60], and Gfi1b [Hoxb8FL; 61], and found that the maxima of all of the ChIP peaks were located at or near the footprints (Fig. 2G,H and Fig. S5E,F). The strong agreement between the Tn5-protected regions and independent evidence of TF binding sites shows that the high depth-of-coverage ATAC-Seq data can be used to track TF binding at the resolution of individual sites. While all the previously known binding sites overlap footprints, the regulatory elements contain additional footprints that do not overlap known binding sites, suggesting a much more complex regulatory scheme than described previously [17, 18, 31, 57, 62].

We determined which TFs might bind to the novel protected regions by scoring their sequences with position weight matrices (PWMs). The protected regions were identified as 5–50bp consecutive nucleotides having less than a threshold number of cuts (Section 4 and Fig. S6–S9). With this criterion, CRMs 7 and 18 had 15 and 21 potential footprints respectively, including previously known binding sites. The sequence of each novel footprint was scored with a set of immune-cell specific TF PWMs from TRANSFAC [63]. 5 footprints matched C/EBP-family, Runx1, Ikzf1, and Jun/Fos on CRM 7 (Fig. S6) while 7 footprints matched Runx1, Ikzf1, MZF1, Jun/Fos, GATA-3, and Sp1 on CRM 18 (Fig. S7). There is a preponderance of matches to Runx1 and C/EBP-family TF sites (Table S2). This type of architecture, where an enhancer has many low-affinity binding sites for a single TF, has been shown to increase enhancer sensitivity through a process known as suboptimization [20, 21]. The large number of sites for the two TFs predicts that CRMs 7 and 18 are highly sensitive to the expression of Runx1 and C/EBP-family TFs.

### 2.4 Accessibility Dynamics and the *cis*-Regulatory Logic of *Cebpa* Enhancers

Having determined the architecture of *Cebpa* regulatory regions at binding site resolution, we next compared the dynamics of accessibility and reporter expression. We computed total accessibility as the sum of Tn5 cuts over each regulatory element during differentiation (Fig. 3). Since enhancers boost gene expression to a level higher than that driven by the promoter itself, we compared their accessibility to the fold change of the enhancer-bearing reporters (Fig. 1F). While *Cebpa* is upregulated two-fold in GCSF conditions (Fig. 1C), the total accessibility of its promoter does not change much, declining slightly during pre-treatment and then recovering after OHT induction. We observed a correlation between enhancer accessibility and fold change for all the enhancers. CRMs 7 and 18 have both higher fold change and higher accessibility in GCSF compared to IL3, whereas both fold change and accessibility of CRM 16 decrease in time (Fig. 3). Despite the correlation, a strict causal link between total accessibility and fold change is lacking. The fold change of CRMs 7 and 18 increases and peaks at earlier time points during GCSF pre-treatment but total accessibility either doesn’t change at all (CRM 18) or increases slightly (CRM 7). Similarly, while the total accessibility of CRMs 7 and 18 peaks 96 hrs after OHT treatment in GCSF conditions, their fold change declines over that interval. Therefore, even though the total accessibility of the two enhancers at 96 hrs is ∼2.5 times its value at -48 hrs, the upregulation relative to the expression of the promoter at 96 hrs is comparable to or lower than its initial value.

**Figure 3:**
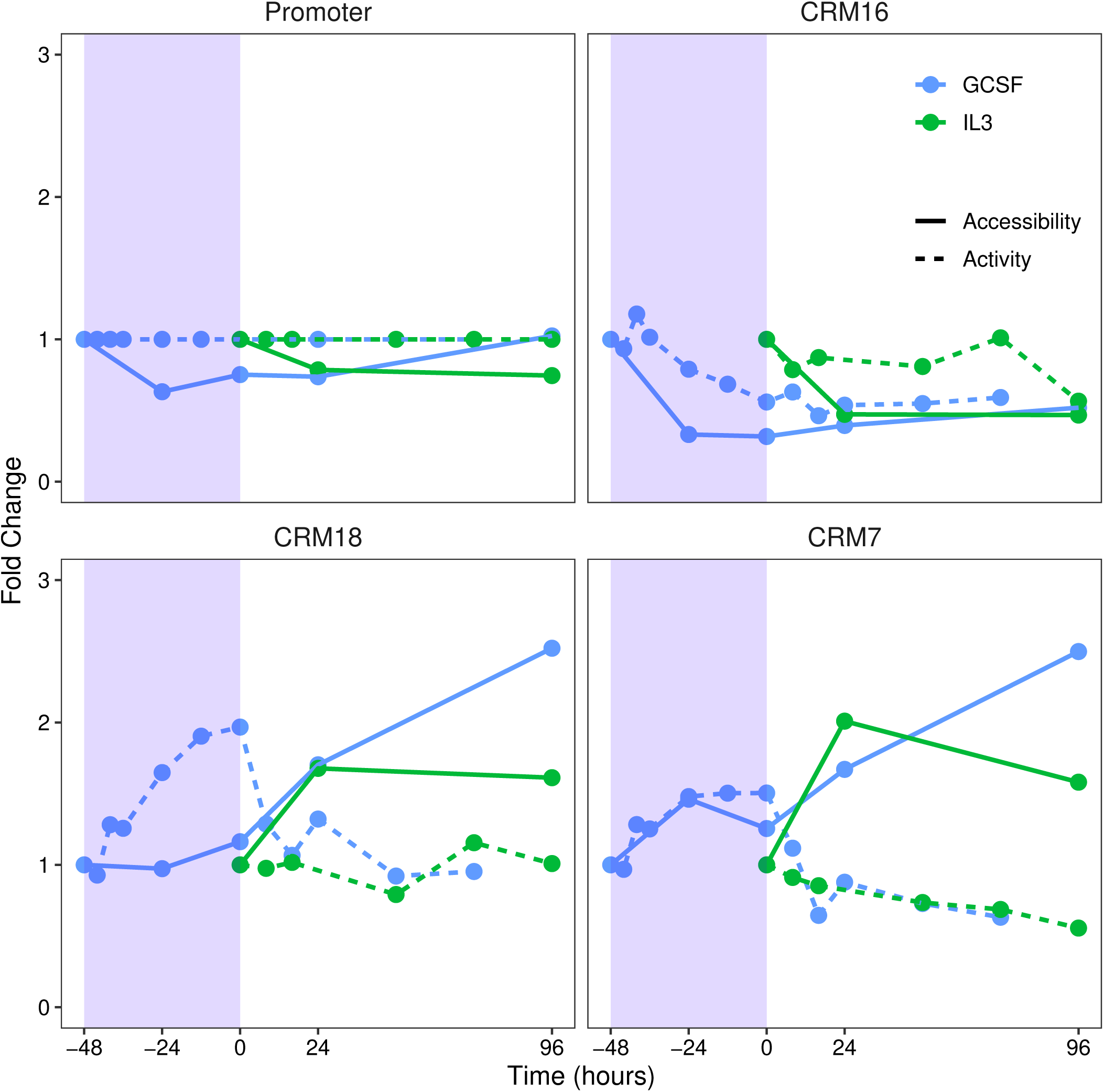
The dynamics of enhancer accessibility and expression. Fold change in the expression of enhancer-bearing reporters relative to promoter (dotted) and total accessibility (solid) are plotted together. Fold change and accessibility were normalized to their values in undifferentiated PUER cells. Differentiation in macrophage (IL3) and neutrophil (GCSF) conditions is shown in green and blue respectively. Blue shaded region shows the period of GCSF pre-treatment prior to induction with OHT during neutrophil differentiation.

The lack of a causal link between total accessibility and fold change led us to examine the accessibility dynamics of individual nucleotides within CRMs 7 and 18. The accessibility of a few positions does, in fact, increase during GCSF pre-treatment (Fig. 4A,C, S10, and S11). We divided the two enhancers uniformly into bins and characterized the accessibility within each bin (Fig. 4B,D). Of 9 bins in CRM 7, the accessibility of only one, bin 5, increases during the pre-treatment period, while all but three respond after OHT treatment. Notably, bin 5 contains binding sites for C/EBP-family TFs. Of 13 bins in CRM 18, bins 3 and 5, also containing C/EBP-family TF sites, increase in accessibility during GCSF pre-treatment, while 9 of 13 are OHT responsive. The change in average accessibility between consecutive timepoints is 6–10 fold higher in the vicinity of C/EBP sites as compared to the remainder of the enhancer (Fig. S12A,B). These data indicate that only the accessibility of nucleotides adjacent to C/EBP binding sites increases during GCSF treatment while OHT treatment results in a broad increase in accessibility across each enhancer.

**Figure 4:**
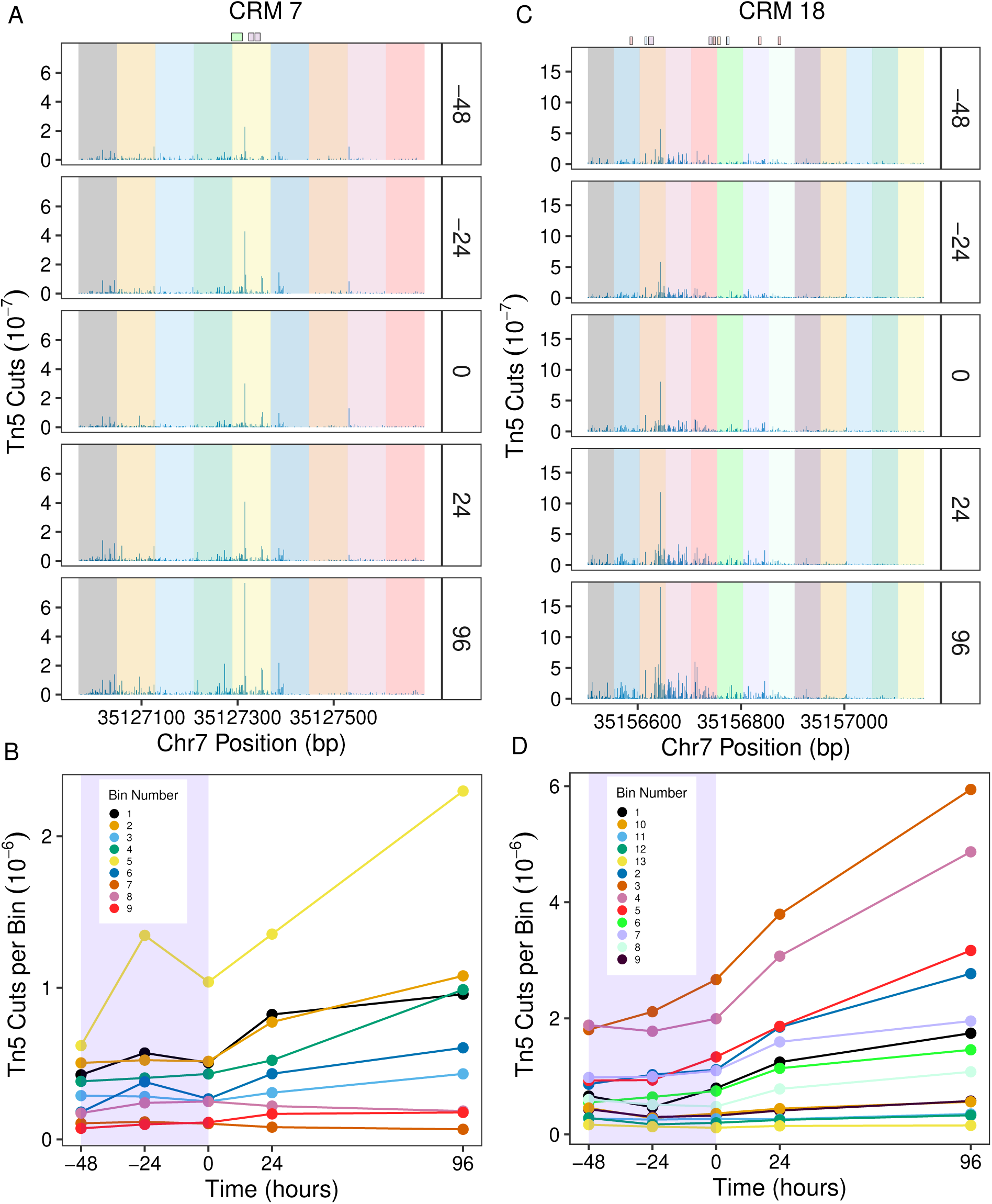
Binding site-resolution dynamics of enhancer accessibility during differentiation. **A,C**. The panels show Tn5 cuts at each nucleotide of CRMs 7 (panel A) and 18 (panel C) at different time points in GCSF conditions. Each enhancer has been uniformly divided into bins shown as shaded regions. TF binding sites are shown at the top; see Figure 2E,F for legend. Replicates have been pooled and normalized by library size. **B,D**. Time series of average accessibility in each bin for CRMs 7 (panel B) and 18 (panel D). Blue shaded region shows period of GCSF pre-treatment prior to induction with OHT during neutrophil differentiation.

The accessibility of binding site-adjacent nucleotides to Tn5 increases with TF occupancy [Fig. S15; 39, 40], implying that the observed increase in accessibility is a result of increased occupancy of C/EBP binding sites. The early increase in accessibility, being localized to a few nucleotides, is a small proportion of the total and therefore does not increase it appreciably. OHT induction, in contrast, results in a broad increase in accessibility across the length of the enhancers (Fig. 4, S10, S11, S13, and S14). The main consequence of OHT induction is the activation of the PUER protein, resulting in an increase in PU.1 binding at its sites (Fig. S15, S16, and 2F,H). The broad increase in accessibility is most likely a result of PU.1’s non-conventional pioneering activity as it is known to be capable of displacing nucleosomes to increase the accessibility of enhancers [35, 36]. These data show that the OHT-dependent increase in the accessibility of the enhancers does not translate into an appreciable increase in the upregulation driven by them (Fig. 3).

## 3 Discussion

The transcription of developmental genes is regulated in a complex scheme, involving multiple enhancers [3, 64–66], and multiple TFs/binding sites per enhancer [16, 17, 67, 68]. This makes it challenging to determine which particular TF or subset of TFs causally govern gene expression in particular tissues or at particular time points during development. We utilized coupled time series datasets of DNA accessibility and gene expression to find causal links between the chromatin state and gene expression of *Cebpa* enhancers.

The comparisons of total accessibility and gene expression did not support a causal link between the two. We computed the fold change in the expression of the reporter bearing both an enhancer and the *Cebpa* promoter to that of a promoter-only reporter as a measure of the activity of the enhancer. Accessibility and fold change do not follow a consistent pattern when comparing enhancers. While CRM 18 has roughly two-fold higher accessibility than CRM 7 in undifferentiated cells (Figs. S10 and S11), its fold change is slightly lower than CRM 7’s (Fig. 1F). There is a lack of correspondence when comparing accessibility and fold change in time as well. Although there is a general correlation between the total accessibility of CRM 7 and 18 and the fold change in reporter expression driven by them, where both properties increase during differentiation, the sequence of events shows that increased accessibility could not be causing greater fold change. The increase in accessibility follows the increase in fold change instead of preceding it (Fig. 3). The discrepancy is particularly striking in the period following PU.1 activation by OHT, during which the accessibility of CRMs 7 & 18 increases ∼2.5- and ∼2-fold in GCSF and IL3 conditions respectively (Fig. 3). Both the CRMs have PU.1 binding sites in their vicinity (Fig. 2F, H and S16) and the increased accessibility is consistent with PU.1’s role as a pioneer TF [35–38]. Despite the broad increase in accessibility across the two enhancers, fold change declines from its peak at 0 hrs, showing that increased accessibility is not sufficient to upregulate enhancer activity.

This result is surprising since DNA accessibility is thought to regulate transcription by limiting the access of TFs to their binding sites [7–12]. A potential reason for the lack of a causal link between total accessibility and fold change is that accessibility is a limiting factor only when it is very low and nucleosome occupancy is high. Once accessibility increases beyond a threshold value, other factors such as TF occupancy and activity, might drive gene expression. In support of this explanation, we observed a coordinate reduction in both the accessibility and fold change of CRM 16, which has a relatively low fold change of 1.17–1.38 during the initial 12 hours of GCSF pre-treatment (Fig. 1F) and could be near the hypothesized accessibility threshold. In contrast, CRMs 7 and 18 have a fold change of ∼2 in undifferentiated PUER cells and are perhaps above the accessibility threshold. For these enhancers, TF occupancy, rather than the overall accessibility of the enhancer, drive the increase in fold change during GCSF pre-treatment (Fig. 4 and S12). To summarize, the apparent disconnect between total accessibility and fold change might be reconciled with a model in which, after an initial opening, the upregulation caused by enhancers is primarily driven by TF occupancy and activity.

Although TF binding sites can be detected in DNase-Seq and ATAC-Seq data [12, 47, 49], there has been mixed success in identifying individual or even aggregate footprints for some TFs [39, 69]. Using the accessibility profiles based on pooled data (Fig. 2E,F and S5C,D) we were able to recover all of the previously known binding sites as individual footprints. Whether this can be generalized to genes other than *Cebpa* will be the subject of future work. However, success in correctly identifying 21 binding sites for about 10 different TFs provides hope that high depth-of-sequencing ATAC-Seq libraries coupled with the bias correction method developed here might improve the genome-wide detection of binding sites. While the *Cebpa* promoter and enhancers were already known to have a complex organization [17, 18, 31, 32], these accessibility profiles reveal many new footprints, corresponding to about 6 TFBSs per element. In particular, CRM 7 is predicted to bind C/EBP at 3 additional sites (Fig. S6) and CRM 18 is predicted to bind Runx1 at 5 new sites (Fig. S7). This composition of the enhancers, with multiple weak sites instead of a few strong sites is reminiscent of “suboptimization” [20, 21], which allows specific and sensitive control of gene expression in response to the TF.

One potential limitation of our analysis is that the reporters are in a heterologous location in the ROSA26 locus that may not reflect the regulation of *Cebpa* elements on chromosome 7. This is mitigated by the observation that when promoters are placed in different genomic environments, they maintain their relative strength relationships despite great variation in absolute expression [70]. Furthermore, integration of the reporters into the same site ensures that the genomic environment is controlled for. Even if there are any genomic location-specific effects, they are not expected to affect the temporal dynamics of fold change. A second limitation is that the enhancers were not placed at the same distance from the promoter as in the endogenous *Cebpa* locus. A recent study utilized the piggyBac transposon to vary the distance between an enhancer and its cognate promoter showed that enhancer activity decreased with distance, with 60%-80% activity at ±40kb [71]. The fold change measurements therefore likely overestimate enhancer strength by virtue of being placed closer than in the endogenous locus, although this effect should be uniform across time points.

Developmental genes have complex regulatory schemes with multiple enhancers, each potentially bound by several TFs at a large number of sites. Although the regulatory logic of a few well-studied loci is well understood, the complexity of regulatory architecture has limited our ability to infer cause-effect relationships. Our results show that profiling chromatin state at high genomic and temporal resolution coupled to reporter data can provide insight into the causality of regulatory events directing differentiation.

## 4 Materials and Methods

### 4.1 PUER Cell Culture and Differentiation Protocol

PUER cells were cultured and differentiated as described previously [18, 28]. Briefly, PUER cells were routinely maintained in complete Iscove’s Modified Dulbecco’s Glutamax medium (IMDM; Gibco, 12440061) supplemented with 10% FBS, 50*µ*M *β*-mercaptoethanol, 5ng/ml IL3 (Peprotech, 213-13). PUER cells were differentiated into macrophages by adding 200nM 4-hydroxy-tamoxifen (OHT; Sigma, H7904-5MG). Cells were differentiated into neutrophils by replacing IL3 with 10ng/ml Granulocyte Colony Stimulating Factor (GCSF; Peprotech, 300-23) and inducing with 100nM OHT after 48 hours.

### 4.2 Reporter knock-in cell line construction

#### 4.2.1 Cas9 and donor vector design and cloning

Cas9 and the sgRNA were expressed from the pSpCas9(BB)-2A-GFP (PX458) plasmid from Addgene [72, #48138;]. The 20-nt guide (5’-ACTGGAGTTGCAGATCACGA-3’) targets a location (113053001–113053020 chr 6; mm10 coordinates) in the first intron of *Gt(ROSA)26Sor*. The guide was cloned into the vector by Gibson Assembly (Gibson Assembly Cloning Kit; New England BioLabs #E5510S) according to the manufacturer’s instructions. The vector was linearized by cutting at two BbsI sites that follow the hU6 promoter. The insert was made by amplifying part of the hU6 promoter with a reverse primer that includes the guide sequence, with an extra G at the 5’ end for enhanced expression, and sequence homologous to the pSpCas9(BB)-2A-GFP vector (Fwd primer: p-1; Rev primer: p-2; Table S1).

The donor vectors were constructed in the pGL4.17[*luc2/Neo*] backbone from Promega (#E672A). DNA homologous to the genomic sequence flanking the Cas9 cleavage site at position 113053017 on chr 6 (“homology arms”) was cloned into the vector. Homology arm 1 (HA1; 113053018–113053656) was amplified from mouse genomic DNA with a forward primer that included the guide RNA target followed by GGG (Fwd primer: p-3; Rev primer: p-4; Table S1). Both primers also contained homology to pGL4.17 for cloning into the SpeI site. HA2 (113052448–113053017) was amplified with a reverse primer that included the guide RNA target followed by GGG (Fwd primer: p-5; Rev primer: p-6; Table S1). HA2 was cloned into the SalI site of the vector. Inclusion of two guide RNA target sites in the donor vector allowed its linearization in cells according to the double-cut donor strategy [41]. The *Cebpa promoter* was cloned between BglII and HindIII (Fwd primer: p-7; Rev primer: p-8; Table S1). CRM 7 was cloned in the BamHI site (Fwd primer: p-9; Rev primer: p-10; Table S1). CRM 18 was cloned in the BstBI site (Fwd primer: p-11; Rev primer: p-12; Table S1). CRM 16 was cloned in the BstBI site (Fwd primer: p-13; Rev primer: p-14; Table S1).

#### 4.2.2 Antibiotic selection, limiting dilution, and screening

200,000 PUER cells were transfected with 500ng each of Cas9 and donor vector in SF buffer (Lonza, V4SC-2096) using program CM134 of the 4D-Nucleofector (Lonza). Cells were allowed to synthesize *Neo* protein product for 72 hours before initiating antibiotic selection. Cells were selected with 1.6 mg/mL G-418 at a density of 2,500 cells/mL for 10 days. Limiting dilution was performed by seeding ∼30 cells per 96-well plate in G-418-free medium. Cells were fed on day 4 and colonies were transferred to a 24-well plate on day 8. On day 14, 200*µ*L of the cell suspension was washed twice in PBS and genomic DNA was extracted with Quickextract solution (Lucigen, QE0905T) according to the manufacturer’s protocol.

Four primers used in three pairs were used to identify the knock-in lines in which the reporter gene had been inserted in a site-specific and biallelic manner. The “out” primers (out-1; out-2; Table S1) flank the homology arms on chromosome 6, 213bp after HA1 and 246bp before HA2 respectively. The “in” primers are complementary to sequences in the insert (in-1; in-2; Table S1). in-1 is complementary to the *Cebpa* promoter and elongates towards out-1, while in-2 is complementary to the *Neo* gene and elongates towards out-2. Amplicons of the predicted size resulting from PCR with in-1/out-1 and in-2/out-2 indicated successful HDR of each junction. PCR with the out-1/out-2 pair has three outcomes. If site-specific integration did not occur, the PCR is expected to produce a 1.7kb amplicon. Biallelic site-specific integration is expected to result in amplicons ranging from ∼5.3kb to ∼7.5kb depending on composition of the insert. Monoallelic site-specific integration is expected to result in both the 1.7kb and the longer amplicon. The in/out PCRs were carried out with Q5 polymerase (New England BioLabs, M0494S), while the out/out PCRs were carried out with PrimeSTAR GXL DNA Polymerase (Takara Bio, R051A). Seamless HDR was confirmed by sequencing the junctions between the insert and the mouse genome. This process generated 2, 5, 3, and 1 independent confirmed biallelic lines for Promoter, CRM 18, CRM 16, and CRM 7 respectively. Clones 9, 31, 7, and 7 were chosen for Promoter, CRM 18, CRM 16, and CRM 7 respectively for further experiments.

### 4.3 Reporter Assays

#### 4.3.1 Bulk Reporter Assays

PUER cells were seeded at a density of 2.5 × 10^5^ cells/mL in 6 replicates before being differentiated as described above. Cells were sampled by washing twice in PBS and lysed in Glo Lysis Buffer (Promega, E2661) for 5 min at RT. The lysate was cleared by centrifugation at 12,000 rpm at 4^◦^C for 5 min. The Luciferase assay was performed with 100 *µ*L of the cleared lysate (Steady Glo reagent, Promega, E2510) according to the manufacturer’s instructions. Total protein concentration was determined with the remaining 100 *µ*L of the lysate using the BCA assay (Thermo Scientific, 23227) and a standard curve constructed with Albumin. Luminescence and absorbance were measured in a multimode plate reader (Beckman Coulter, DTX-880). Cell number was also estimated using the CellTox Green Cytotoxicity Assay (Promega, G8743) according to manufacturer’s instructions.

#### 4.3.2 Single-cell Reporter Assays

Single-cell flow cytometry-based Luciferase measurements were done by intracellular immunostaining of Firefly Luciferase. 0.5–1 million cells of each reporter gene bearing line were sampled at each time point of the differentiation and spiked with ∼0.25 million cells of the Promoter line stained with CFSE (eBioscience, 65-0850). The CFSE-stained cells can be distinguished in the FITC channel during flow cytometry and their Luciferase signal provides an internal standard. In addition to reporter lines, PUER cells lacking Luciferase were also sampled to estimate background non-specific staining during flow cytometry. The spiked samples were stained with the LIVE/DEAD Fixable Violet Dead Cell Stain Kit (Invitrogen, L34955) to exclude dead cells. Cell were fixed and permeabilized with BD Cytofix/Cytoperm Fixation and Permeabilization Solution (BD Biosciences, 554722) according to the manufacturer’s protocol. Cells were then stained for intracellular Firefly Luciferase with anti-Firefly Luciferase antibody (clone EPR17789, Abcam, ab185923) at a final concentration of 2.37 *µ*g/mL (1:75 dilution) and polyclonal Goat Anti-Rabbit IgG H&L conjugated to Alexa Fluor 647 (Abcam, ab150083) at a final concentration of 1 *µ*g/mL (1:2000 dilution). Fluorescence was recorded on a BD FACSymphony analyzer.

Luciferase expression, in units of undifferentiated Promoter expression, was calculated as follows. Let *f*_sample_ and 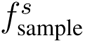 be the median fluorescence of the stained Luciferase protein in the sample and the spiked Promoter internal standard respectively. Also, let *f*_PUER_ and 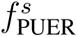 be the median fluorescence in the PUER cells without Luciferase and Promoter internal standard they were spiked with respectively. The background non-specific fluorescence in the sample was estimated as

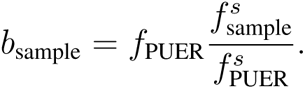

The Luciferase expression in the sample was then computed as

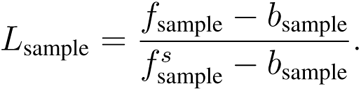

### 4.4 Fast-ATAC

Fast-ATAC was conducted as described by Corces *et al.* [50]. 50,000 cells/sample were collected and centrifuged at 500g for 5 minutes at 4^◦^C. The supernatant was removed and the resulting pellet was resuspended in a 50µL transposition mixture consisting of 25µL 2x TD buffer, 2.5µL TDE1 (Illumina, FC-121-1030), 0.5µL 1% digitonin (Promega, G9441), and 22µL nuclease-free water. The resuspended cells were then incubated at 37^◦^C with shaking at 300 rpm for 30 minutes. DNA was purified using a Qiagen MinElute Reaction Cleanup Kit (Qiagen, 28204) and eluted in 10µL of elution buffer (10mM Tris-HCl, pH 8).

The transposed DNA library was amplified for a limited number of cycles as described by Buenrostro *et al.* [12]. First, 10µL of transposed DNA was amplified in a 50*µ*L PCR reaction using NEBNext High-Fidelity 2x PCR Master Mix (New England BioLabs, M0541S) for 5 cycles after a 5 min elongation step. Primers included adaptors for sequencing and were barcoded according to Illumina guidelines for pooling. The PCR reaction was held at 4^◦^C while 5*µ*L of its product was amplified in a 15*µ*L qPCR reaction with NEBNext High-Fidelity Master Mix and SYBR Green I (Invitrogen, S7563) at a final concentration of 0.5x. The number of PCR cycles needed to reach 1/4 of the maximum fluorescence was determined and the 50*µ*L PCR reaction was continued for as many additional cycles (9–11). The PCR product was purified using a Qiagen MinElute PCR Purification Kit and eluted in 20µL elution buffer (10mM Tris-HCl, pH 8). The libraries were then analyzed on an Agilent Bioanalyzer for QC before being pooled in equimolar amounts. The pooled libraries were sequenced in the 2 × 150bp format to an average depth of 300 million reads/sample on a NovaSeq 6000 S4 (Illumina) by Psomagen Inc.

### 4.5 Data Processing and analysis

Raw sequence files were tested for quality using FastQC (v. 0.11.5). Reads were trimmed to remove adaptor sequences and filtered to have a minimum phred score of 33 and a minimum length of 20bp using Trimmomatic (v 0.39). Trimmomatic was run with the command-line options `-PE -phred33 ILLUMINACLIP:<TRIMMOMATIC-HOME>/adapters/NexteraPE-PE.fa:2:30:10:3:TRUE MINLEN:20`, where `<TRIMMOMATIC-HOME>` is the path to the Trimmomatic directory. The reads were aligned to the *mm10* genome, allowing for inserts up to 2kb in length, using Hisat2 (v. 2.2.0). Hisat2 was run with the command-line options `-x -no-spliced-alignment -I 10 -X 2000`. Duplicates were marked using the `MarkDuplicates` tool of Picard (v. 2.23.1) package. Duplicate reads were removed during the read counting process in R.

#### 4.5.1 Read Counting

Tn5 cuts were counted in R (v. 4.1.0) using a custom script where each read was counted as one Tn5 cut. Properly paired and unique reads were read using the GenomicAlignments package [73] with filtering parameters `isPaired = T, isProperPair = T, isUnmappedQuery = F, isSecondaryAlignment = F, isNotPassingQualityControls = F, isDuplicate = F, isSupplementaryAlignment = F`. Tn5 leaves a 9bp 5’ overhang [Fig. S3A; 12, 52], so that the 5’ ends of a pair of reads resulting from the same transposition event map to different positions on the + and - strands. The read mapping to the + strand starts on the 1st nucleotide of the overhang whereas the one mapping to the - strand starts on the 9th nucleotide of the overhang. It is common to shift the Tn5 cut position by +4/-5 bp for reads mapping to the +/- strand respectively [12]. This however does not have the desired effect of assigning the same position to the two reads resulting from the same transposition event (Fig. S3B). For both cuts to have the same location (5th nucleotide of the overhang), they must shifted by +4/-4 bp (Fig. S3B,C). Finally, Tn5 cuts in each sample were normalized to their respective library size.

#### 4.5.2 Bias Correction

Tn5 transposase does not bind to and cut naked DNA uniformly but exhibits bias for certain motifs [39, 40, 46, 52]. We tested two previously published Tn5 sequence bias correction methods, HINT-ATAC [52] and BaG-Foot [39], as well as a new method we developed in this study. Tn5 DNA sequence preferences were inferred from ATAC-Seq libraries prepared by transposing purified mouse genomic DNA with Tn5 (GSM2981009 [53], GSM1550786 [54], GSM4048700 [55], and GSM2333650/GSM2333651 [56]).

##### Kmer-based bias inference

Tn5 binding preferences were inferred from purified DNA ATAC-Seq libraries by estimating the probability of Tn5 cutting at each possible hexamer. Since hexamers occur at different frequencies in the genome, the bias *b*(*w_k_*) at hexamer *w_k_*, where *k* = 1*, …*, 4096, was computed as the ratio of the frequency of observed Tn5 cuts *p_c_*(*w_k_*) to the frequency of the occurrence of the hexamer in the mouse genome *p*(*w_k_*),

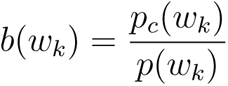

All the purified DNA ATAC-Seq datasets were merged into one BAM file and the MakeBiasCorrectionTableBAM function of the BagFoot package [39] was used with parameters np = 6, atac = T to compute the Tn5 bias at each hexamer.

##### HINT-ATAC bias correction

This was performed as described [52]. The number of bias-corrected cuts *x_i_* at position *i* were computed as

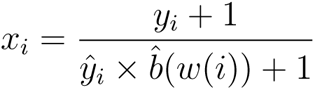

where *y_i_* are the observed cuts, *w*(*i*) is the hexamer at *i*, and *b*(*w*) is the Tn5 bias at hexamer 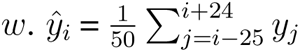 is the mean of observed cuts in a 50bp window around 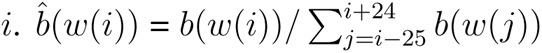 is the Tn5 bias at position *i* relative to the total bias over the 50bp window (Fig. S4, second panel).

##### BaGFoot bias correction

This was performed as described [39]. The bias-corrected cuts *x_i_* at position *i* were computed as

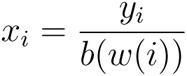

where *y_i_* are the observed cuts at position *i* and *b*(*w*(*i*)) is the bias of the hexamer *w*(*i*) at position *i* (Fig. S4, third panel).

##### Our bias correction method

HINT-ATAC corrects the observed cuts with a factor that depends on the bias at the position relative to average bias in a 50bp window and the average number of cuts in the window. As a result, it does not correct fine-scale bias at the resolution of individual binding sites and also adds one cut to each position even if it had no cuts originally (Fig. S4). While BaGFoot does not average the cuts or the bias, it can amplify noise at low-accessibility nucleotides which could mask protected regions.

We developed a method that corrects bias at the resolution of individual nucleotides without amplifying noise at positions with low counts. The method adjusts the cuts at individual nucleotides but only for positions that have more cuts than the background level. We modeled the cuts at each position with the Poisson distribution 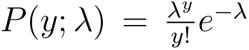, where *λ* is the mean number of cuts in the sample across the genome. We correct the bias for a position *i* if 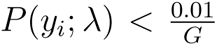, where *G* is the size of the mouse genome (Bonferroni correction) so that the corrected cuts are

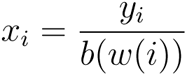

where *b*(*w*(*i*)) is the bias of the hexamer at *i* (Fig. S4, fourth panel).

### 4.6 Identification of new binding sites in *Cebpa* enhancers

Protected regions were identified as runs of 5–50bp consecutive low-accessibility nucleotides bordered by positions having more cuts than an enhancer-specific threshold. Cuts were pooled over all samples for this analysis. The thresholds were 400, 100, 40, or 200 in the promoter, CRM 7, 16, or 18 respectively. Putative TFs binding to the protected regions were identified using TRANSFAC’s *Match* tool [74, https://gene-regulation.com] using immune-specific position weight matrices (PWMs).

## Acknowledgments

We thank M. Takaku, S. Nechaev, and A. Dhasarathy for discussions and comments. We thank the UND Genomics core for services. This work was supported by the National Science Foundation [1942471 to M.] and the National Institute of General Medical Sciences of the National Institutes of Health under Award number 5P20GM104360.

## Author contributions

T. E. L. designed and conducted experiments, wrote software, analyzed data, and wrote the manuscript. T. B. designed and conducted experiments, analyzed data, and wrote the manuscript. A. R. designed and conducted experiments. M. N. designed and conducted experiments. S. N. designed and conducted experiments, analyzed data, and wrote the manuscript. S. K. designed and conducted experiments. M. conceptualized and supervised the work, designed experiments, wrote software, analyzed data, and wrote the manuscript.

## Materials & correspondence

Requests for materials and correspondence should be addressed to M.

## References

[1] Spitz F, Furlong EEM. Transcription factors: from enhancer binding to developmental control. Nat Rev Genet. 2012 Sep;13(9):613–26.

[2] May G, Soneji S, Tipping AJ, Teles J, McGowan SJ, Wu M, et al. Dynamic analysis of gene expression and genome-wide transcription factor binding during lineage specification of multipotent progenitors. Cell Stem Cell. 2013 Dec;13(6):754–68.

[3] Choi J, Lysakovskaia K, Stik G, Demel C, Söding J, Tian TV, et al. Evidence for additive and synergistic action of mammalian enhancers during cell fate determination. Elife. 2021 03;10.

[4] Mouse ENCODE Consortium, Stamatoyannopoulos JA, Snyder M, Hardison R, Ren B, Gingeras T, et al. An encyclopedia of mouse DNA elements (Mouse ENCODE). Genome Biol. 2012;13(8):418.

[5] Kagey MH, Newman JJ, Bilodeau S, Zhan Y, Orlando DA, van Berkum NL, et al. Mediator and cohesin connect gene expression and chromatin architecture. Nature. 2010 Sep;467(7314):430–5.

[6] Boija A, Klein IA, Sabari BR, Dall’Agnese A, Coffey EL, Zamudio AV, et al. Transcription Factors Activate Genes through the Phase-Separation Capacity of Their Activation Domains. Cell. 2018 Dec;175(7):1842– 1855.e16.

[7] Kornberg RD, Lorch Y. Chromatin structure and transcription. Annu Rev Cell Biol. 1992;8:563–87.

[8] Bartsch J, Truss M, Bode J, Beato M. Moderate increase in histone acetylation activates the mouse mammary tumor virus promoter and remodels its nucleosome structure. Proc Natl Acad Sci U S A. 1996 Oct;93(20):10741–6.

[9] Mellor J. The dynamics of chromatin remodeling at promoters. Mol Cell. 2005 Jul;19(2):147–57.

[10] Boyle AP, Davis S, Shulha HP, Meltzer P, Margulies EH, Weng Z, et al. High-resolution mapping and characterization of open chromatin across the genome. Cell. 2008 Jan;132(2):311–22.

[11] Thurman RE, Rynes E, Humbert R, Vierstra J, Maurano MT, Haugen E, et al. The accessible chromatin landscape of the human genome. Nature. 2012 Sep;489(7414):75–82.

[12] Buenrostro JD, Giresi PG, Zaba LC, Chang HY, Greenleaf WJ. Transposition of native chromatin for fast and sensitive epigenomic profiling of open chromatin, DNA-binding proteins and nucleosome position. Nat Methods. 2013 Dec;10(12):1213–8. Cited in PubMed; PMID 24097267.

[13] Ptashne M. Epigenetics: core misconcept. Proc Natl Acad Sci U S A. 2013 Apr;110(18):7101–3.

[14] Boyes J, Felsenfeld G. Tissue-specific factors additively increase the probability of the all-or-none formation of a hypersensitive site. EMBO J. 1996 May;15(10):2496–507.

[15] Adams CC, Workman JL. Binding of disparate transcriptional activators to nucleosomal DNA is inherently cooperative. Mol Cell Biol. 1995 Mar;15(3):1405–21.

[16] Kim AR, Martinez C, Ionides J, Ramos AF, Ludwig MZ, Ogawa N, et al. Rearrangements of 2.5 kilobases of noncoding DNA from the Drosophila even-skipped locus define predictive rules of genomic cis-regulatory logic. PLoS Genet. 2013 Feb;9(2):e1003243.

[17] Cooper S, Guo H, Friedman AD. The +37 kb Cebpa Enhancer Is Critical for Cebpa Myeloid Gene Expression and Contains Functional Sites that Bind SCL, GATA2, C/EBP, PU.1, and Additional Ets Factors. PLoS One. 2015;10(5):e0126385.

[18] Repele A, Krueger S, Bhattacharyya T, Tuineau MY, Manu. The regulatory control of Cebpa enhancers and silencers in the myeloid and red-blood cell lineages. PLoS One. 2019;14(6):e0217580.

[19] Levine M, Davidson EH. Gene regulatory networks for development. Proc Natl Acad Sci U S A. 2005 Apr;102(14):4936–42.

[20] Kim DS, Risca VI, Reynolds DL, Chappell J, Rubin AJ, Jung N, et al. The dynamic, combinatorial cisregulatory lexicon of epidermal differentiation.;53:1564–1576.

[21] Farley EK, Olson KM, Zhang W, Brandt AJ, Rokhsar DS, Levine MS. Suboptimization of developmental enhancers.; 350:325–328.

[22] Zhang DE, Zhang P, Wang ND, Hetherington CJ, Darlington GJ, Tenen DG. Absence of granulocyte colonystimulating factor signaling and neutrophil development in CCAAT enhancer binding protein alpha-deficient mice. Proc Natl Acad Sci U S A. 1997 Jan;94(2):569–74.

[23] Legraverend C, Antonson P, Flodby P, Xanthopoulos KG. High level activity of the mouse CCAAT/enhancer binding protein (C/EBP alpha) gene promoter involves autoregulation and several ubiquitous transcription factors. Nucleic Acids Res. 1993 Apr;21(8):1735–42.

[24] Rosen ED, Hsu CH, Wang X, Sakai S, Freeman MW, Gonzalez FJ, et al. C/EBPalpha induces adipogenesis through PPARgamma: a unified pathway. Genes Dev. 2002 Jan;16(1):22–6.

[25] Wu C, Jin X, Tsueng G, Afrasiabi C, Su AI. BioGPS: building your own mash-up of gene annotations and expression profiles. Nucleic Acids Res. 2016 Jan;44(D1):D313–6.

[26] Lattin JE, Schroder K, Su AI, Walker JR, Zhang J, Wiltshire T, et al. Expression analysis of G Protein-Coupled Receptors in mouse macrophages. Immunome Res. 2008 Apr;4:5.

[27] Scott EW, Simon MC, Anastasi J, Singh H. Requirement of transcription factor PU.1 in the development of multiple hematopoietic lineages. Science. 1994 Sep;265(5178):1573–7.

[28] Dahl R, Walsh JC, Lancki D, Laslo P, Iyer SR, Singh H, et al. Regulation of macrophage and neutrophil cell fates by the PU.1:C/EBPalpha ratio and granulocyte colony-stimulating factor. Nat Immunol. 2003 Oct;4(10):1029–36.

[29] Gonzalez D, Luyten A, Bartholdy B, Zhou Q, Kardosova M, Ebralidze A, et al. ZNF143 protein is an important regulator of the myeloid transcription factor C/EBP. J Biol Chem. 2017 Nov;292(46):18924– 18936.

[30] Guo H, Cooper S, Friedman AD. In Vivo Deletion of the Cebpa +37 kb Enhancer Markedly Reduces Cebpa mRNA in Myeloid Progenitors but Not in Non-Hematopoietic Tissues to Impair Granulopoiesis. PLoS One. 2016;11(3):e0150809.

[31] Bertolino E, Reinitz J, Manu. The analysis of novel distal Cebpa enhancers and silencers using a transcriptional model reveals the complex regulatory logic of hematopoietic lineage specification. Dev Biol. 2016 May;413(1):128–44.

[32] Peng L, Guo H, Ma P, Sun Y, Dennison L, Aplan PD, et al. HoxA9 binds and represses the Cebpa +8 kb enhancer. PLoS One. 2019;14(5):e0217604.

[33] Walsh JC, DeKoter RP, Lee HJ, Smith ED, Lancki DW, Gurish MF, et al. Cooperative and antagonistic interplay between PU.1 and GATA-2 in the specification of myeloid cell fates. Immunity. 2002 Nov;17(5):665–76.

[34] Laslo P, Spooner CJ, Warmflash A, Lancki DW, Lee HJ, Sciammas R, et al. Multilineage transcriptional priming and determination of alternate hematopoietic cell fates. Cell. 2006 Aug;126(4):755–66.

[35] Barozzi I, Simonatto M, Bonifacio S, Yang L, Rohs R, Ghisletti S, et al. Coregulation of transcription factor binding and nucleosome occupancy through DNA features of mammalian enhancers. Mol Cell. 2014 Jun;54(5):844–857.

[36] Minderjahn J, Schmidt A, Fuchs A, Schill R, Raithel J, Babina M, et al. Mechanisms governing the pioneering and redistribution capabilities of the non-classical pioneer PU.1. Nat Commun. 2020 Jan;11(1):402.

[37] Ghisletti S, Barozzi I, Mietton F, Polletti S, De Santa F, Venturini E, et al. Identification and characterization of enhancers controlling the inflammatory gene expression program in macrophages. Immunity. 2010 Mar;32(3):317–28.

[38] Feng R, Desbordes SC, Xie H, Tillo ES, Pixley F, Stanley ER, et al. PU.1 and C/EBPalpha/beta convert fibroblasts into macrophage-like cells. Proc Natl Acad Sci U S A. 2008 Apr;105(16):6057–62.

[39] Baek S, Goldstein I, Hager GL. Bivariate Genomic Footprinting Detects Changes in Transcription Factor Activity. Cell Rep. 2017 05;19(8):1710–1722.

[40] Bentsen M, Goymann P, Schultheis H, Klee K, Petrova A, Wiegandt R, et al. ATAC-seq footprinting unravels kinetics of transcription factor binding during zygotic genome activation. Nat Commun. 2020 08;11(1):4267.

[41] Zhang JP, Li XL, Li GH, Chen W, Arakaki C, Botimer GD, et al. Efficient precise knockin with a double cut HDR donor after CRISPR/Cas9-mediated double-stranded DNA cleavage. Genome Biol. 2017 Feb;18(1):35.

[42] Klemm SL, Shipony Z, Greenleaf WJ. Chromatin accessibility and the regulatory epigenome.; 20:207–220.

[43] Coux RX, Owens NDL, Navarro P. Chromatin accessibility and transcription factor binding through the perspective of mitosis.; 11:236–240.

[44] Mirny LA. Nucleosome-mediated cooperativity between transcription factors. Proc Natl Acad Sci U S A. 2010 Dec;107(52):22534–9.

[45] Anderson JD, Thåström A, Widom J. Spontaneous access of proteins to buried nucleosomal DNA target sites occurs via a mechanism that is distinct from nucleosome translocation. Mol Cell Biol. 2002 Oct;22(20):7147–57.

[46] Karabacak Calviello A, Hirsekorn A, Wurmus R, Yusuf D, Ohler U. Reproducible inference of transcription factor footprints in ATAC-seq and DNase-seq datasets using protocol-specific bias modeling. Genome Biol. 2019 02;20(1):42.

[47] Gusmao EG, Dieterich C, Zenke M, Costa IG. Detection of active transcription factor binding sites with the combination of DNase hypersensitivity and histone modifications. Bioinformatics. 2014 Nov;30(22):3143– 51.

[48] Yardımcı GG, Frank CL, Crawford GE, Ohler U. Explicit DNase sequence bias modeling enables highresolution transcription factor footprint detection. Nucleic Acids Res. 2014 Oct;42(19):11865–78.

[49] Gusmao EG, Allhoff M, Zenke M, Costa IG. Analysis of computational footprinting methods for DNase sequencing experiments. Nat Methods. 2016 Apr;13(4):303–9.

[50] Corces MR, Buenrostro JD, Wu B, Greenside PG, Chan SM, Koenig JL, et al. Lineage-specific and single-cell chromatin accessibility charts human hematopoiesis and leukemia evolution. Nat Genet. 2016 10;48(10):1193–203.

[51] Sato S, Arimura Y, Kujirai T, Harada A, Maehara K, Nogami J, et al. Biochemical analysis of nucleosome targeting by Tn5 transposase. Open Biol. 2019 08;9(8):190116.

[52] Li Z, Schulz MH, Look T, Begemann M, Zenke M, Costa IG. Identification of transcription factor binding sites using ATAC-seq. Genome Biol. 2019 02;20(1):45.

[53] Senft AD, Costello I, King HW, Mould AW, Bikoff EK, Robertson EJ. Combinatorial Smad2/3 Activities Downstream of Nodal Signaling Maintain Embryonic/Extra-Embryonic Cell Identities during Lineage Priming. Cell Rep. 2018 08;24(8):1977–1985.e7.

[54] Pastor WA, Stroud H, Nee K, Liu W, Pezic D, Manakov S, et al. MORC1 represses transposable elements in the mouse male germline. Nat Commun. 2014 Dec;5:5795.

[55] Onimaru K, Tatsumi K, Tanegashima C, Kadota M, Nishimura O, Kuraku S. Developmental hourglass and heterochronic shifts in fin and limb development. Elife. 2021 02;10.

[56] Gray LT, Yao Z, Nguyen TN, Kim TK, Zeng H, Tasic B. Layer-specific chromatin accessibility landscapes reveal regulatory networks in adult mouse visual cortex. Elife. 2017 01;6.

[57] Guo H, Ma O, Speck NA, Friedman AD. Runx1 deletion or dominant inhibition reduces Cebpa transcription via conserved promoter and distal enhancer sites to favor monopoiesis over granulopoiesis. Blood. 2012 May;119(19):4408–18.

[58] Heinz S, Benner C, Spann N, Bertolino E, Lin YC, Laslo P, et al. Simple combinations of lineage-determining transcription factors prime cis-regulatory elements required for macrophage and B cell identities. Mol Cell. 2010 May;38(4):576–89.

[59] Garber M, Yosef N, Goren A, Raychowdhury R, Thielke A, Guttman M, et al. A high-throughput chromatin immunoprecipitation approach reveals principles of dynamic gene regulation in mammals. Mol Cell. 2012 Sep;47(5):810–22.

[60] Zhao L, Glazov EA, Pattabiraman DR, Al-Owaidi F, Zhang P, Brown MA, et al. Integrated genome-wide chromatin occupancy and expression analyses identify key myeloid pro-differentiation transcription factors repressed by Myb. Nucleic Acids Res. 2011 Jun;39(11):4664–79.

[61] Nestorowa S, Hamey FK, Pijuan Sala B, Diamanti E, Shepherd M, Laurenti E, et al. A single-cell resolution map of mouse hematopoietic stem and progenitor cell differentiation. Blood. 2016 08;128(8):e20–31.

[62] Guo H, Ma O, Friedman AD. The Cebpa +37-kb enhancer directs transgene expression to myeloid progenitors and to long-term hematopoietic stem cells. J Leukoc Biol. 2014 Sep;96(3):419–26.

[63] Matys V, Kel-Margoulis OV, Fricke E, Liebich I, Land S, Barre-Dirrie A, et al. TRANSFAC and its module TRANSCompel: transcriptional gene regulation in eukaryotes. Nucleic Acids Res. 2006 Jan;34(Database issue):D108–10.

[64] Whyte WA, Orlando DA, Hnisz D, Abraham BJ, Lin CY, Kagey MH, et al. Master transcription factors and mediator establish super-enhancers at key cell identity genes. Cell. 2013 Apr;153(2):307–19.

[65] Marinić M, Aktas T, Ruf S, Spitz F. An Integrated Holo-Enhancer Unit Defines Tissue and Gene Specificity of the Fgf8 Regulatory Landscape. Developmental Cell. 2013 12;(0):–. Available from: http://www.sciencedirect.com/science/article/pii/S1534580713000737.

[66] Hong JW, Hendrix DA, Levine MS. Shadow enhancers as a source of evolutionary novelty. Science. 2008 Sep;321(5894):1314.

[67] Wilson NK, Timms RT, Kinston SJ, Cheng YH, Oram SH, Landry JR, et al. Gfi1 expression is controlled by five distinct regulatory regions spread over 100 kilobases, with Scl/Tal1, Gata2, PU.1, Erg, Meis1, and Runx1 acting as upstream regulators in early hematopoietic cells. Mol Cell Biol. 2010 Aug;30(15):3853–63.

[68] Kittler R, Zhou J, Hua S, Ma L, Liu Y, Pendleton E, et al. A comprehensive nuclear receptor network for breast cancer cells. Cell Rep. 2013 Feb;3(2):538–51.

[69] D’Oliveira Albanus R, Kyono Y, Hensley J, Varshney A, Orchard P, Kitzman JO, et al. Chromatin information content landscapes inform transcription factor and DNA interactions. Nat Commun. 2021 02;12(1):1307.

[70] Hong CKY, Cohen BA. Genomic environments scale the activities of diverse core promoters. Genome Res. 2022 Jan;32(1):85–96.

[71] Zuin J, Roth G, Zhan Y, Cramard J, Redolfi J, Piskadlo E, et al. Nonlinear control of transcription through enhancer-promoter interactions. Nature. 2022 04;604(7906):571–577.

[72] Ran FA, Hsu PD, Wright J, Agarwala V, Scott DA, Zhang F. Genome engineering using the CRISPR-Cas9 system. Nat Protoc. 2013 Nov;8(11):2281–308.

[73] Lawrence M, Huber W, Pagès H, Aboyoun P, Carlson M, Gentleman R, et al. Software for computing and annotating genomic ranges. PLoS Comput Biol. 2013;9(8):e1003118.

[74] Kel AE, Gössling E, Reuter I, Cheremushkin E, Kel-Margoulis OV, Wingender E. MATCH: A tool for searching transcription factor binding sites in DNA sequences. Nucleic Acids Res. 2003 Jul;31(13):3576–9.

