## Supplementary Information for "The contributions of DNA accessibility and transcription factor occupancy to enhancer activity during cellular differentiation"

Trevor Long[†], Tapas Bhattacharyya[†], Andrea Repele, Madison Naylor, Sunil Nooti, Shawn Krueger, & Manu[*]

Department of Biology,
University of North Dakota,
10 Cornell Street, Stop 9019, Grand Forks, ND 58202-9019, U.S.A.

[†]Equal contribution
[*]Corresponding author

### S1 Supplementary Information Captions

**Fig S1. Comparison of Luciferase activity in independent lines generated by CRISPR/Cas9 HDR.** Luciferase activity for undifferentiated cells in all confirmed biallelic lines. The luminescence was normalized to cell number estimated using the CellTox assay. In the boxplots, the horizontal line indicates median over 8 replicate measurements, the bottom and top edges are the first the third quartiles, while the vertical lines indicate 1.5 times the interquartile range. Individual replicate measurements are shown as points.

**Fig. S2. Reporter expression and fold change measured by flow cytometry. A.** Dot-plot of CFSE and Alexa Fluor 647 (Luciferase secondary) fluorescence showing the $\mathrm{CFSE}^{+}$ spiked internal control (undifferentiated Promoter) and the $\mathrm{CFSE}^{-}$ sample (CRM 7 in this example). Cells have been gated to exclude debris, doublets/clusters, and dead cells. **B.** Histograms of Luciferase fluorescence intensity of undifferentiated cells of the PUER unedited founder line ($\mathrm{Luc}^{-}$), Promoter, CRM 18, CRM 16, and CRM 7. **C.** Median Luciferase expression in units of $\mathrm{CFSE}^{-}$ spiked undifferentiated promoter during differentiation in GCSF conditions. CRM 18 and Promoter were sampled at high frequency during the first 24 hrs of GCSF pre-treatment and show strong upregulation consistent with bulk Luciferase data (Fig. 1E). **D.** Fold change in median expression relative to the expression of the Promoter, showing upregulation of CRMs 7 and 18 during GCSF pre-treatment.

**Fig S3. Comparison of different strategies for counting Tn5 cuts. A.** Schematic of cleavage by the Tn5 transposase. Adaptor 1 and 2 are Illumina read 1 and 2 adaptors respectively. The pair of green strips represents double stranded genomic DNA. Black triangles are sites of cleavage and integration of the adaptors, showing the 9bp overhang. Sequenced inserts resulting from one Tn5 cleavage event or independent Tn5 cleavages at the same location map to two different positions separated by 8bp depending on whether they align to the + or - strands. The same Tn5 transposition event is counted at two different locations if unmodified left- or right-most mapping positions are used without modification. **B.** Schematic of two different strategies for "centering" the Tn5 cut position. Shifting the + and - strands by +4 and -5 bp respectively [12] does not align the positions correctly, while shifting by +4/-4 bp results in proper centering. **C.** The pattern of Tn5 cuts in CRM 7, pooled over all samples, resulting from the two strategies of shifting a cut's position to the center of the 9bp overhang.

**Fig. S4. Comparison of different strategies for correcting Tn5 DNA sequence bias.** The pattern of Tn5 cuts in CRM 7 in one replicate of the 96h IL3+OHT time point. From top to bottom, bias from Tn5 DNA sequence preferences was either not corrected (Uncorrected) or corrected using HINT-ATAC [52], BagFoot [39], or our method (Section 4).

**Fig. S5. Enhancer-level accessibility profiles of the *Cebpa* promoter and CRM 16. A,B.** Pearson correlation coefficients of Tn5 counts in each regulatory element, averaged with a 10bp sliding window, computed between each pair of samples. **C,D.** Pooled single-nucleotide accessibility profiles. Black bars are the sum of Tn5 counts over all the samples. Red line is a 3bp sliding-window average. Shaded regions show empirically verified known binding sites [18, 23, 31]. **E.** Scaled ChIP-Seq signal of C/EBP$\beta$ in PUER cells (GSM538010). **F.** Scaled ChIP-seq signal of PU.1 (GSM538004) in PUER cells. **A,C,E.** *Cebpa* promoter. **B,D,F.** CRM 16.

**Fig. S6. Novel candidate binding sites in CRM 7.** Pooled single-nucleotide accessibility profiles. Black bars are the sum of Tn5 counts over all the samples. Shaded regions are footprints that match the binding motifs of the indicated TFs (Section 4).

**Fig. S7. Novel candidate binding sites in CRM 18.** Pooled single-nucleotide accessibility profiles. Black bars are the sum of Tn5 counts over all the samples. Shaded regions are footprints that match the binding motifs of the indicated TFs (Section 4).

**Fig. S8. Novel candidate binding sites in the *Cebpa* promoter.** Pooled single-nucleotide accessibility profiles. Black bars are the sum of Tn5 counts over all the samples. Shaded regions are footprints that match the binding motifs of the indicated TFs (Section 4).

**Fig. S9. Novel candidate binding sites in CRM 16.** Pooled single-nucleotide accessibility profiles. Black bars are the sum of Tn5 counts over all the samples. Shaded regions are footprints that match the binding motifs of the indicated TFs (Section 4).

**Fig S10. Single nucleotide-resolution dynamics of CRM 7 accessibility during differentiation.** The panels show Tn5 cuts at each nucleotide of CRM 7 at different time points and conditions. Differentiation in macrophage (IL3) and neutrophil (GCSF) conditions is shown in green and blue respectively. Replicates have been pooled and normalized by library size. The 72bp region bordering the Gfi1 site and containing the C/EBP sites is highlighted with the star.

**Fig. S11. Single nucleotide-resolution dynamics of CRM 18 accessibility during differentiation.** The panels show Tn5 cuts at each nucleotide of CRM 18 at different time points and conditions. Differentiation in macrophage (IL3) and neutrophil (GCSF) conditions is shown in green and blue respectively. Replicates have been pooled and normalized by library size. The 30 bp region bordering the PU.1 and C/EBP sites is highlighted with the star.

**Fig. S12. Comparison of accessibility dynamics of key TF binding sites with the rest of the enhancer.** The temporal change in average accessibility per nucleotide is plotted for key binding sites (red) and for the rest of the enhancer (blue). **A,C.** CRM 7. The red curve is the change in average accessibility over a 72bp region next to the Gfi1 site and spanning C/EBP sites (highlighted in Fig. S10). **B,D.** CRM 18. The red curve is the change in average accessibility over a 30bp region spanning PU.1 and C/EBP sites (highlighted in Fig. S11). **A,B.** Differentiation in neutrophil conditions (GCSF). **C,D.** Differentiation in macrophage conditions (IL3). Blue shaded region shows period of GCSF pre-treatment prior to induction with OHT during neutrophil differentiation.

**Fig. S13. Single nucleotide-resolution dynamics of *Cebpa* promoter accessibility during differentiation.** The panels show Tn5 cuts at each nucleotide of the *Cebpa* promoter at different time points and conditions. Differentiation in macrophage (IL3) and neutrophil (GCSF) conditions is shown in green and blue respectively. Replicates have been pooled and normalized by library size.

**Fig. S14. Single nucleotide-resolution dynamics of CRM 16 accessibility during differentiation.** The panels show Tn5 cuts at each nucleotide of CRM 16 at different time points and conditions. Differentiation in macrophage (IL3) and neutrophil (GCSF) conditions is shown in green and blue respectively. Replicates have been pooled and normalized by library size.

**Fig. S15. The accessibility of nucleotides neighboring PU.1 binding sites increases with PU.1 occupancy.** Genome-wide aggregate PU.1 footprint analysis in undifferentiated PUER cells and 96 hrs after OHT treatment in GCSF conditions. Cut-probability (accessibility) of nucleotides $\pm$100bp from the *Spi1* (PU.1) motif (JASPAR, 2018; MA0080.3). Signal was measured at sites having PWM scores in the top 5%. Undifferentiated PUER cells lack active PU.1 protein and the occupancy of its sites is low. The average accessibility of nucleotides adjacent to the motif is low. OHT treatment activates the PU.1-ER fusion protein, causing it to bind to its sites. The average accessibility of motif-adjacent nucleotides increases significantly.

**Fig. S16. PU.1 binding in the vicinity of CRM 7.** Pooled single-nucleotide accessibility profile of a ±500bp region surrounding CRM 7 in the upper panel. Black bars are the sum of Tn5 counts over all the samples. Shaded regions show empirically verified known binding sites. Scaled ChIP-Seq signal of C/EBP$\beta$ (GSM538010) and PU.1 (GSM538004) in PUER cells are shown in the bottom panel. ChIP signal has been normalized to its maximum in the depicted region.

**Table S1. Primer sequences used in vector construction and screening.**

**Table S2. Chromosomal locations of preliminary novel transcription factor binding sites.**

### S2 Supplementary Tables

| Notation | Sequence |
|---|---|
| out-1 | 5’-CGTCTCGTCGCTGATTGGCTTCTT-3’ |
| out-2 | 5’-GGAGAGAGGCATTCATGGGAGTG-3’ |
| in-1 | 5’-CGGCTGTGGGTAGGAGTTAGAT-3 |
| in-2 | 5’-ATCGCCTTCTACCGCTTGCTCGA-3’ |
| p-1 | 5’-GTACAAAATACGTGACGTAGAAAG-3’ |
| p-2 | 5’-TTATTTTAACTTGCTATTTCTAGCTCTAAAACTCGTGATCTGCAACTCCAGTCGGTGTTTCGTCCTTTCCACAAGATATATA-3’ |
| p-3 | 5’-TCAATATTATTGAAGCATTTATCAGGGTTACTAGTACTGGAGTTGCAGATCACGAGGGAGGGAGGGTCAGCGAAAGT-3’ |
| p-4 | 5’-CCTTAATATTACTTACTTATCCTTGAGAGACGTACTAGTCGAGGGAAGAGGGGGAAG-3’ |
| p-5 | 5’-TGTGTTGGTTTTTTGTGTGTTCGAACTAGATGCTGTCGACTGATCTGCAACTCCAGTCTTTC-3’ |
| p-6 | 5’-GAAGGCTCTCAAGGGCATCGGTCGACACTGGAGTTGCAGATCACGAGGGCTGTCTGGTTTCATGAGTCATC-3’ |
| p-7 | 5’-CCGGTACCTGAGCTCGCTAGCCTCGAGGATATCAGAATTCCCATGGAGATCTAACTCCTACCCACAGCCG-3’ |
| p-8 | 5’-ATTCGCGACCCGAAGCTGAAGCTTCCATGGGAATTCGGCAATCCGGTACTGTTGGTAAAGCCACCATGGA-3’ |
| p-9 | 5’-AGTAAAACCTCTACAAATGTGGTAAAATCGATAAGAATTCCCATGGGGATCCCCCACTTCCACCCCCTAAGA-3’ |
| p-10 | 5’-TGTTAAGGTTGCTCTGCTCAGGGGATCCCCATGGGAATTCGTTTGCGTATTGGGCGCTCTTCCGCTGATCTGCG-3’ |
| p-11 | 5’-TCTTTATTTTCATTACATCTGTGTGTTGGTTTTTTGTGTGGAATTCCCATGGTTCGAAATGCCACCCCTCTGATTTTGC-3’ |
| p-12 | 5’-TCTATTCCAGGGGACTCAGTGTTCGAACCATGGGAATTCCTAGATGCTGTCGACTGATCTGCAACTCCAGTCTTTCTAG-3’ |
| p-13 | 5’-TCTTTATTTTCATTACATCTGTGTGTTGGTTTTTTGTGTGGAATTCCCATGGTTCGAAAAAATCAGTTTATCCCTATGCTGCC-3’ |
| p-14 | 5’-CAGAGTTGTCCTCAGCCCATTCGAACCATGGGAATTCCTAGATGCTGTCGACTGATCTGCAACTCCAGTCTTTCTAG-3’ |

Table S1

| Gene | CRM | Chromosome | Start | End | Tfs |
|---|---|---|---|---|---|
| Cebpa | 7 | 7 | 35127019 | 35127034 | Cebp |
| Cebpa | 7 | 7 | 35127036 | 35127044 | Runx1 |
| Cebpa | 7 | 7 | 35127059 | 35127096 | Cebp, Ikzf1 |
| Cebpa | 7 | 7 | 35127096 | 35127116 | Cebp |
| Cebpa | 7 | 7 | 35127217 | 35127258 | Jun/Fos, Cebp |
| Cebpa | 18 | 7 | 35156512 | 35156526 | MZF1, Runx1 |
| Cebpa | 18 | 7 | 35156578 | 35156593 | Runx1 |
| Cebpa | 18 | 7 | 35156695 | 35156709 | Jun/Fos |
| Cebpa | 18 | 7 | 35156719 | 35156730 | Runx1 |
| Cebpa | 18 | 7 | 35156782 | 35156814 | Runx1, GATA-3, MZF1, Sp1, Ikzf1 |
| Cebpa | 18 | 7 | 35156815 | 35156842 | Jun/Fos, Cebp, Runx1 |
| Cebpa | 18 | 7 | 35156846 | 35156862 | Ikzf1, Runx1 |
| Cebpa | 0 | 7 | 35118286 | 35118321 | MZF1 |
| Cebpa | 0 | 7 | 35118321 | 35118336 | Sp1, MZF1 |
| Cebpa | 0 | 7 | 35118762 | 35118787 | Cebp |
| Cebpa | 0 | 7 | 35118787 | 35118808 | Ikzf1 |
| Cebpa | 0 | 7 | 35118981 | 35119020 | Sp1, MZF1, Runx1 |
| Cebpa | 16 | 7 | 35150786 | 35150805 | Ikzf1, Sp1 |
| Cebpa | 16 | 7 | 35150857 | 35150882 | Runx1 |
| Cebpa | 16 | 7 | 35150899 | 35150933 | Ikzf1, MZF1, Runx1, PU.1 |
| Cebpa | 16 | 7 | 35151025 | 35151042 | PU.1 |
| Cebpa | 16 | 7 | 35151147 | 35151166 | Ikzf1, Runx1 |
| Cebpa | 16 | 7 | 35151335 | 35151355 | Gata3 |
| Cebpa | 16 | 7 | 35151360 | 35151381 | Ikzf1, Runx1 |
| Cebpa | 16 | 7 | 35151381 | 35151411 | Jun/Fos |
| Cebpa | 16 | 7 | 35151411 | 35151441 | Cebp, MZF1, Ikzf1, Runx1 |

Table S2

#### S3 Supplementary Figures

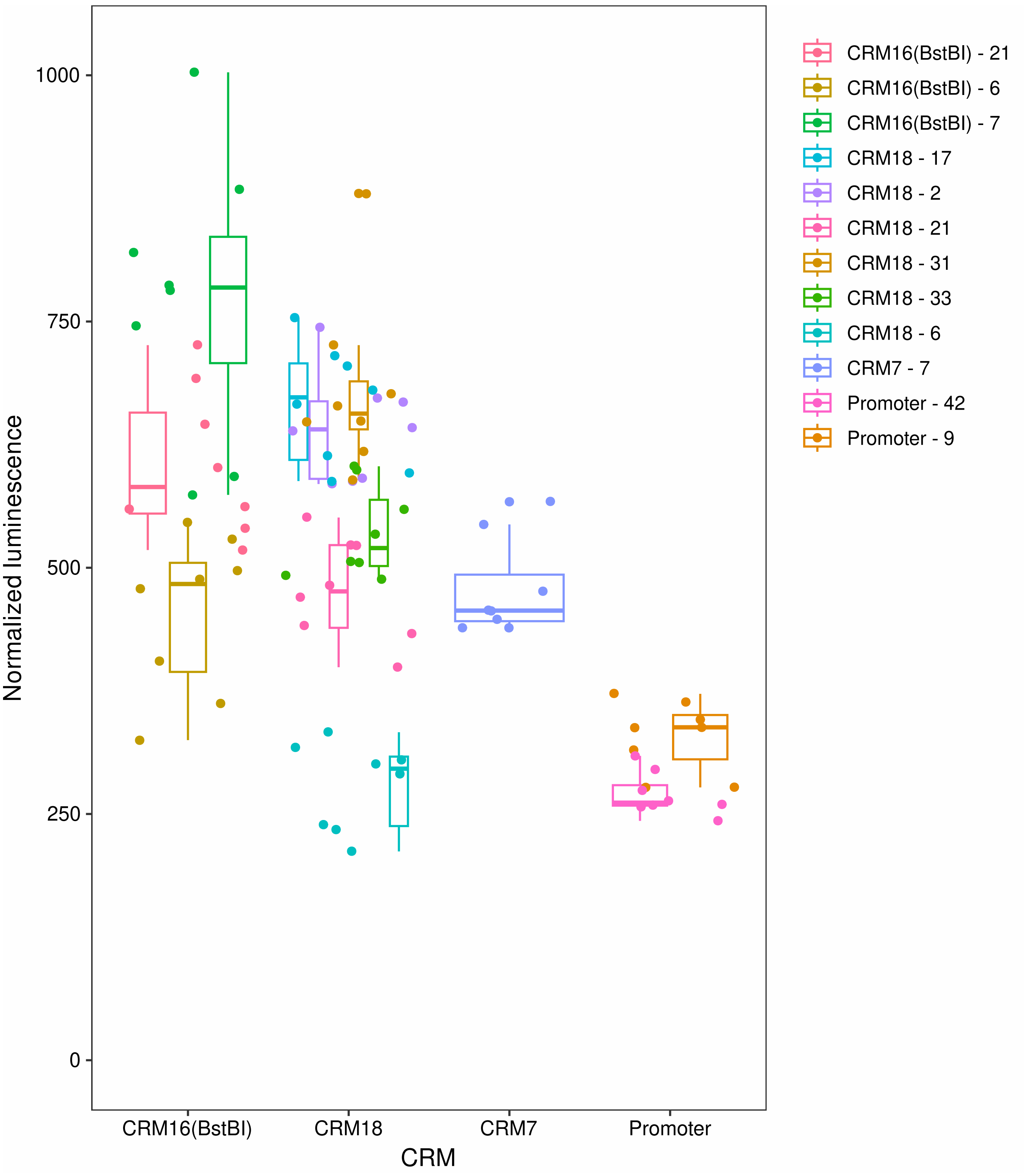


Figure S1

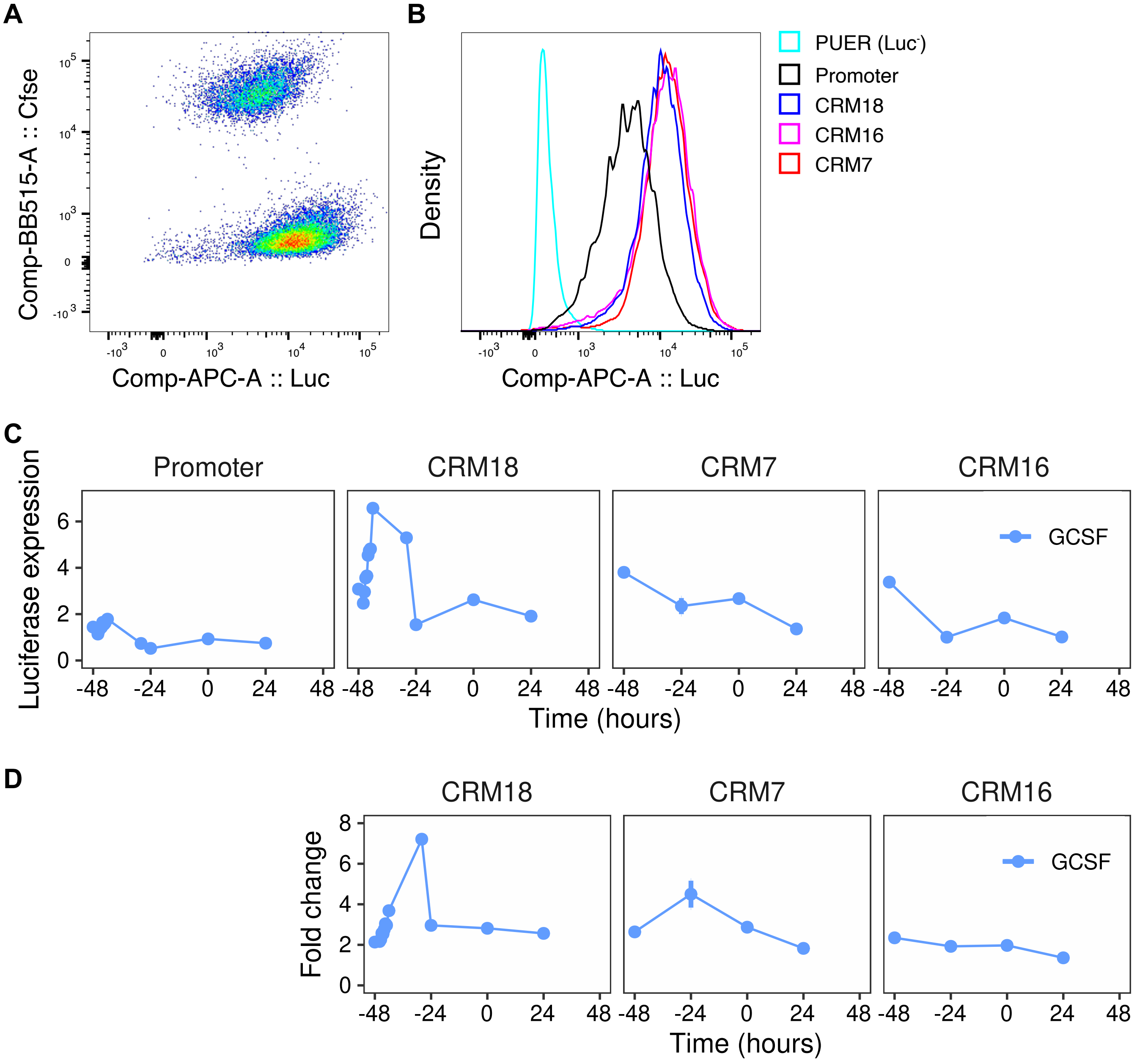
A
B
Comp-BB515-A :: Cfse
Comp-APC-A :: Luc
Density
PUER (Luc-)
Promoter
CRM18
CRM16
CRM7
C
Promoter
CRM18
CRM7
CRM16
Luciferase expression
GCSF
Time (hours)
D
Fold change
Time (hours)

Figure S2

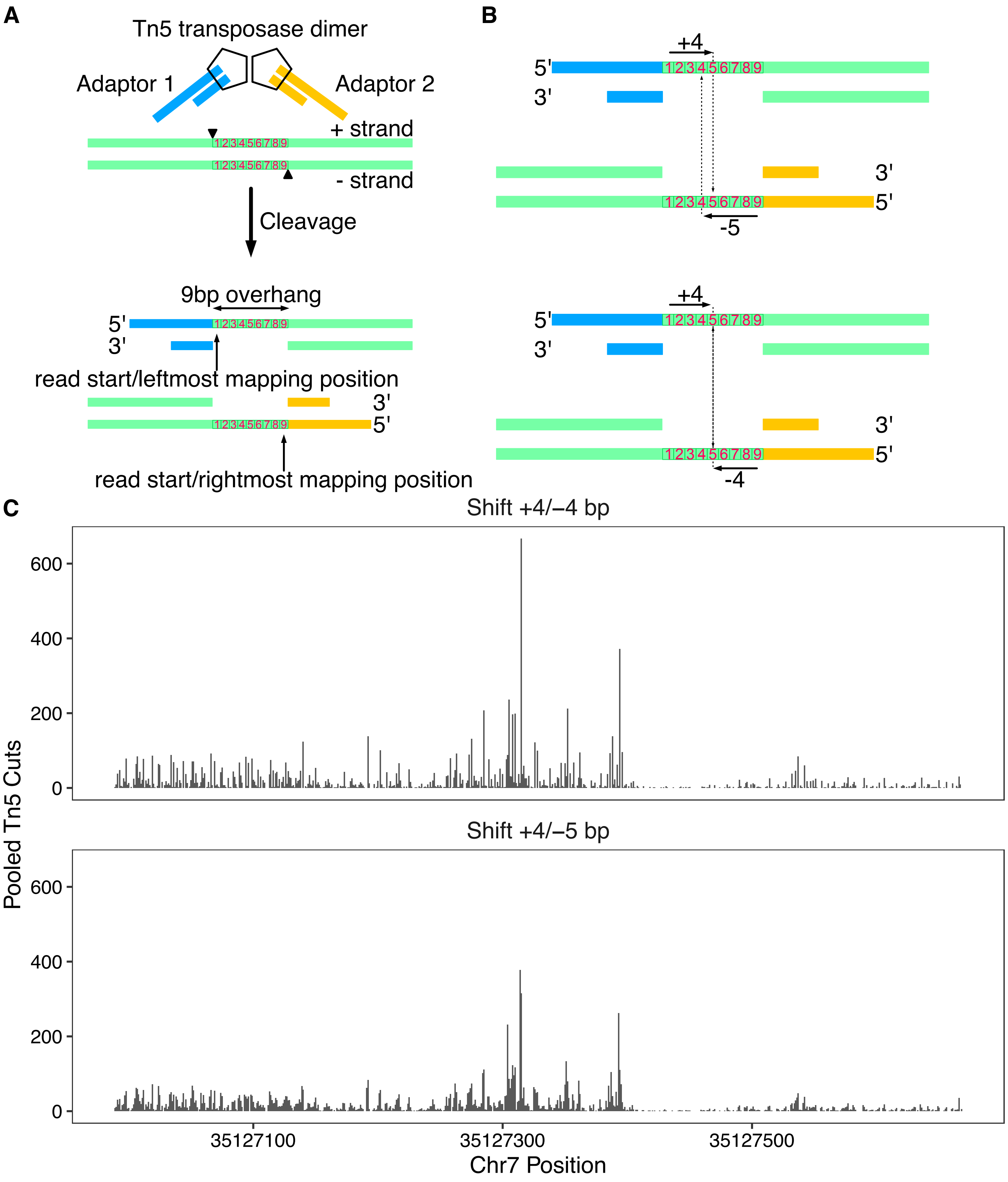
A
Tn5 transposase dimer
Adaptor 1
Adaptor 2
+ strand
- strand
Cleavage
9bp overhang
read start/leftmost mapping position
read start/rightmost mapping position
B
+4
-5
-4
C
Shift +4/–4 bp
Shift +4/–5 bp
Pooled Tn5 Cuts
Chr7 Position
35127100
35127300
35127500

Figure S3

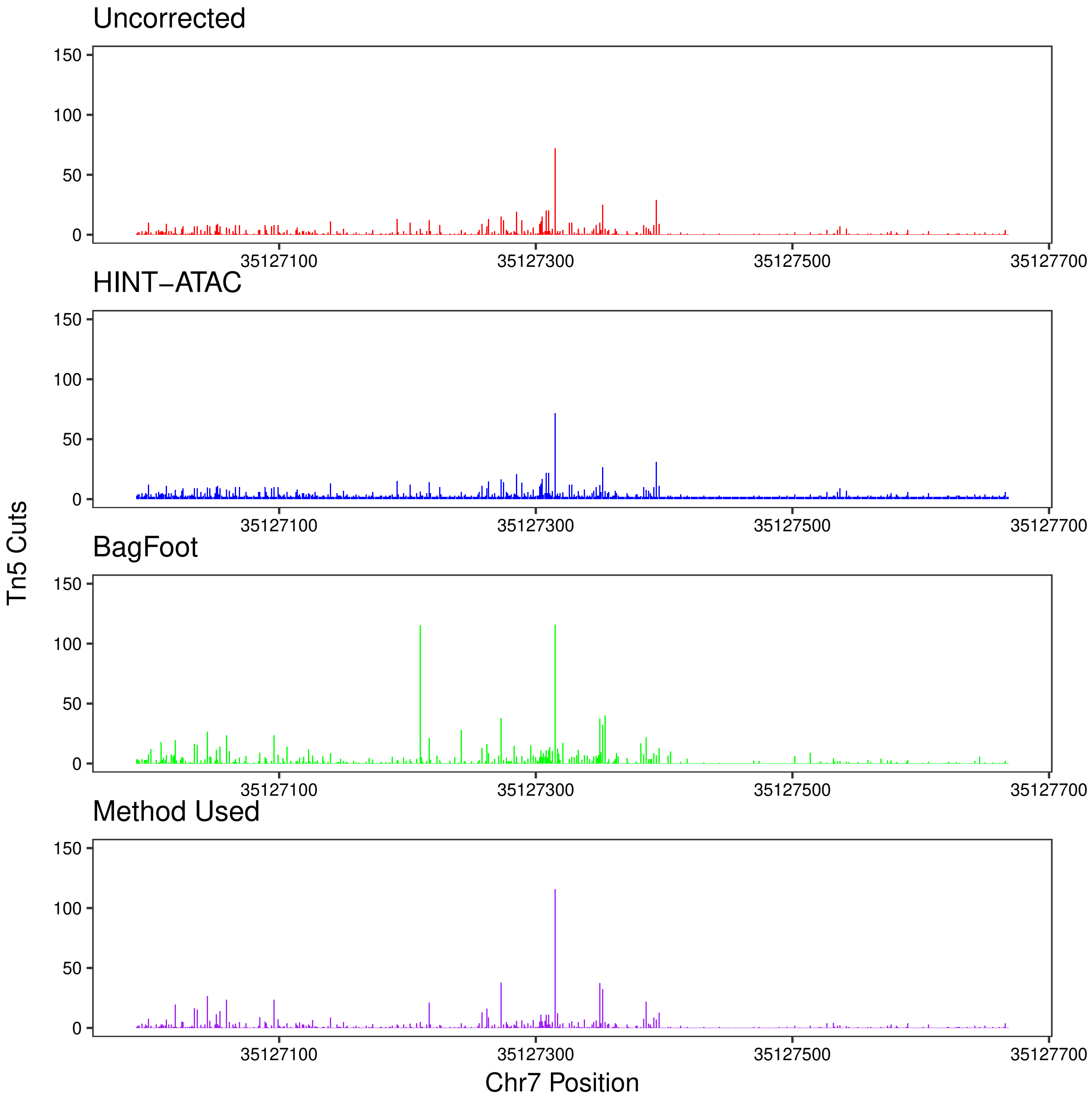

Uncorrected
HINT−ATAC
BagFoot
Method Used
150
100
50
0
35127100
35127300
35127500
35127700
Tn5 Cuts
Chr7 Position


Figure S4

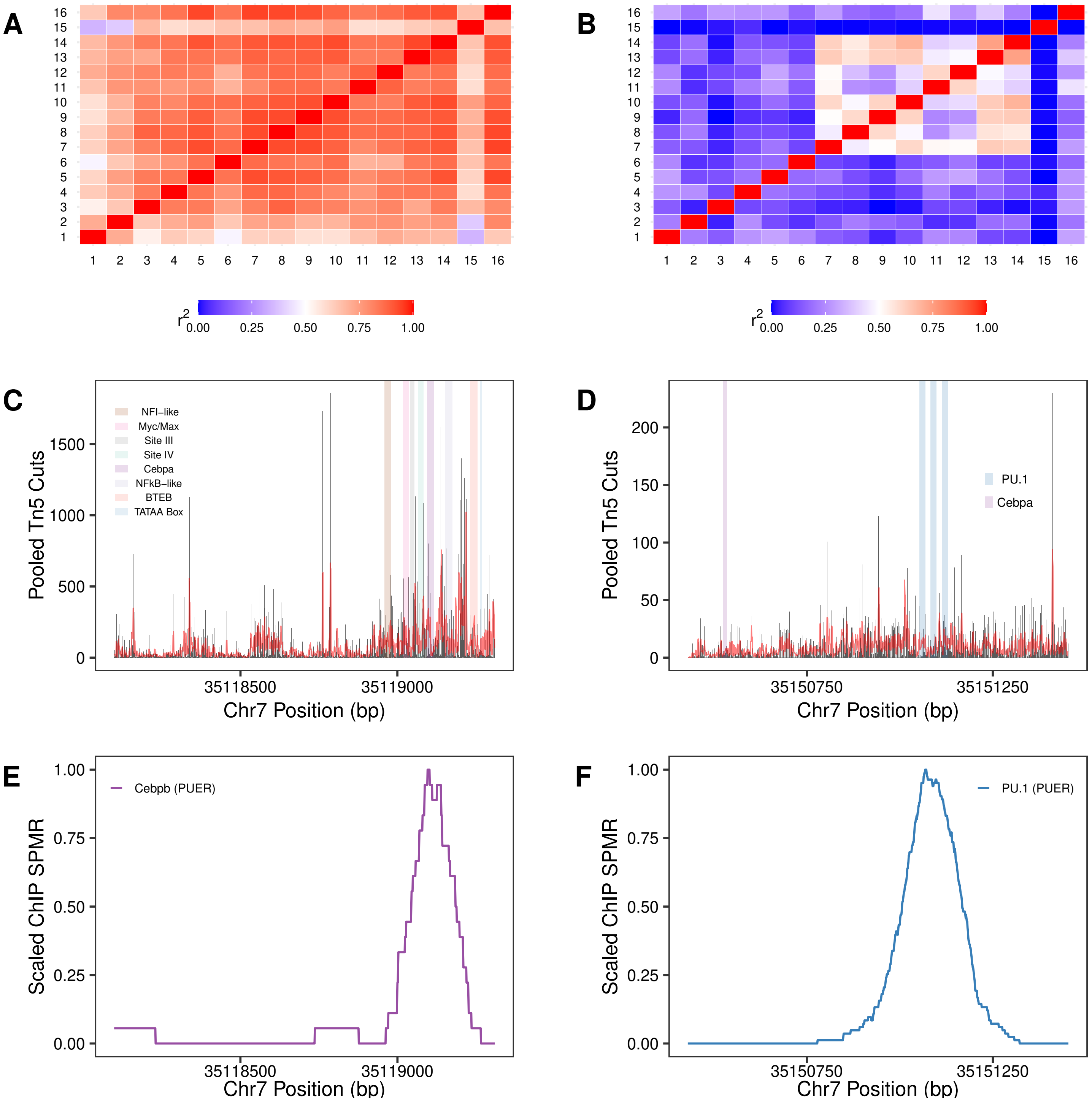
A
B
r2
0.00 0.25 0.50 0.75 1.00
C
Pooled Tn5 Cuts
NFI–like
Myc/Max
Site III
Site IV
Cebpa
NFkB–like
BTEB
TATAA Box
35118500 35119000
Chr7 Position (bp)
D
PU.1
Cebpa
35150750 35151250
E
Scaled ChIP SPMR
Cebpb (PUER)
F
PU.1 (PUER)

Figure S5

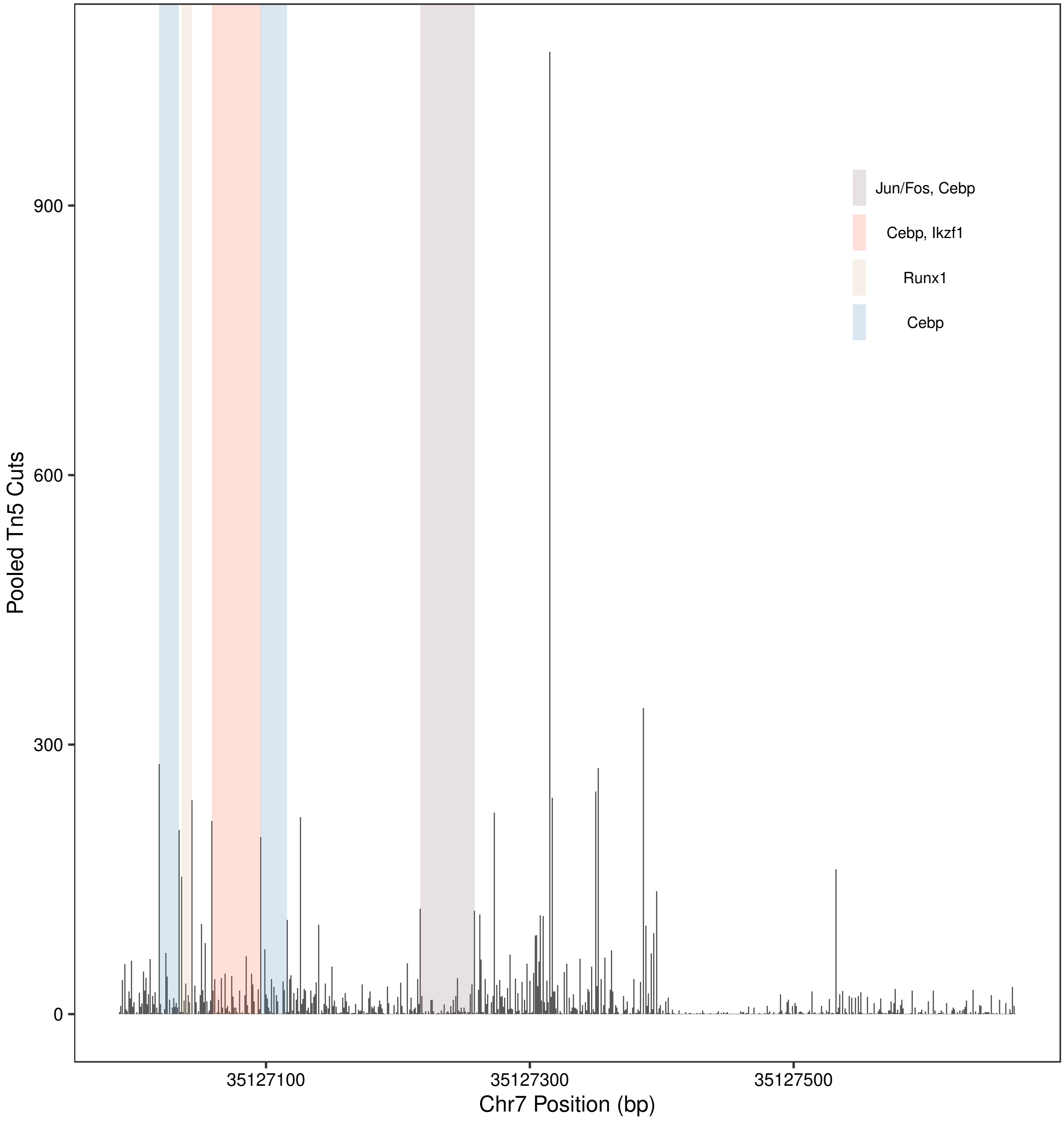
Jun/Fos, Cebp
Cebp, Ikzf1
Runx1
Cebp
Pooled Tn5 Cuts
900
600
300
0
35127100
35127300
35127500
Chr7 Position (bp)

Figure S6

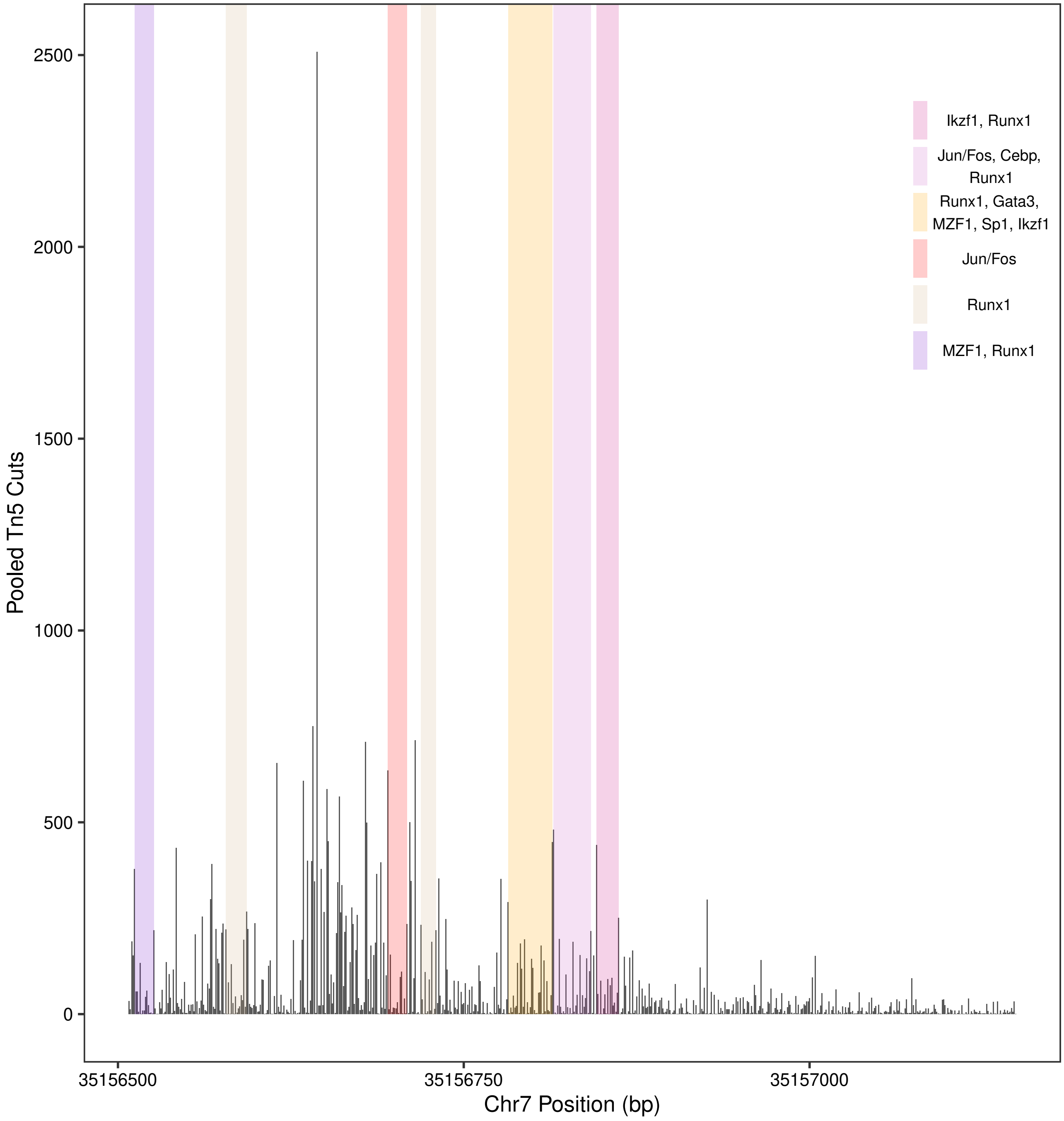

Pooled Tn5 Cuts
2500
2000
1500
1000
500
0
35156500
35156750
35157000
Chr7 Position (bp)
Ikzf1, Runx1
Jun/Fos, Cebp, Runx1
Runx1, Gata3, MZF1, Sp1, Ikzf1
Jun/Fos
Runx1
MZF1, Runx1


Figure S7

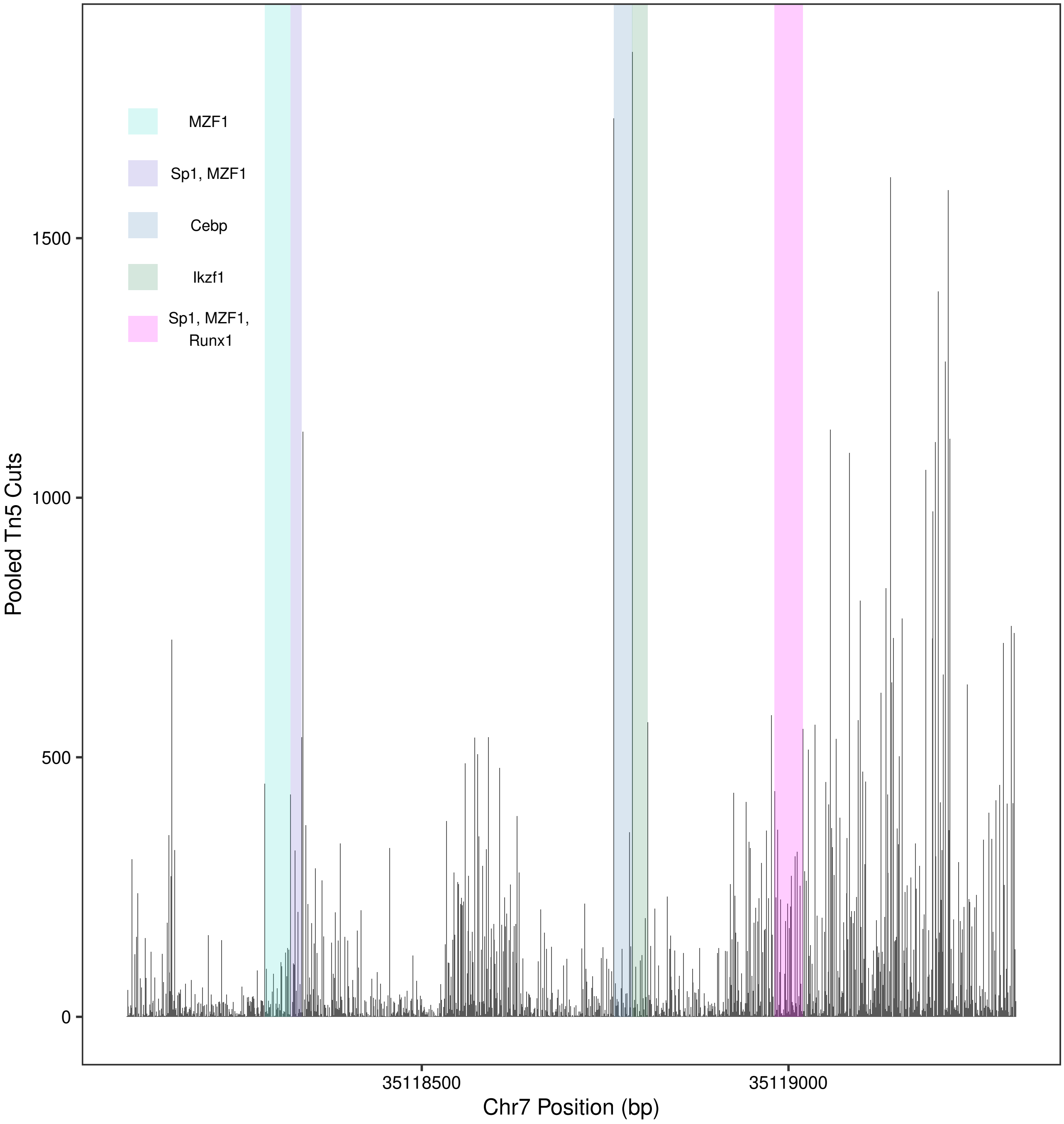
MZF1
Sp1, MZF1
Cebp
Ikzf1
Sp1, MZF1, Runx1
1500
1000
500
0
Pooled Tn5 Cuts
35118500
35119000
Chr7 Position (bp)

Figure S8

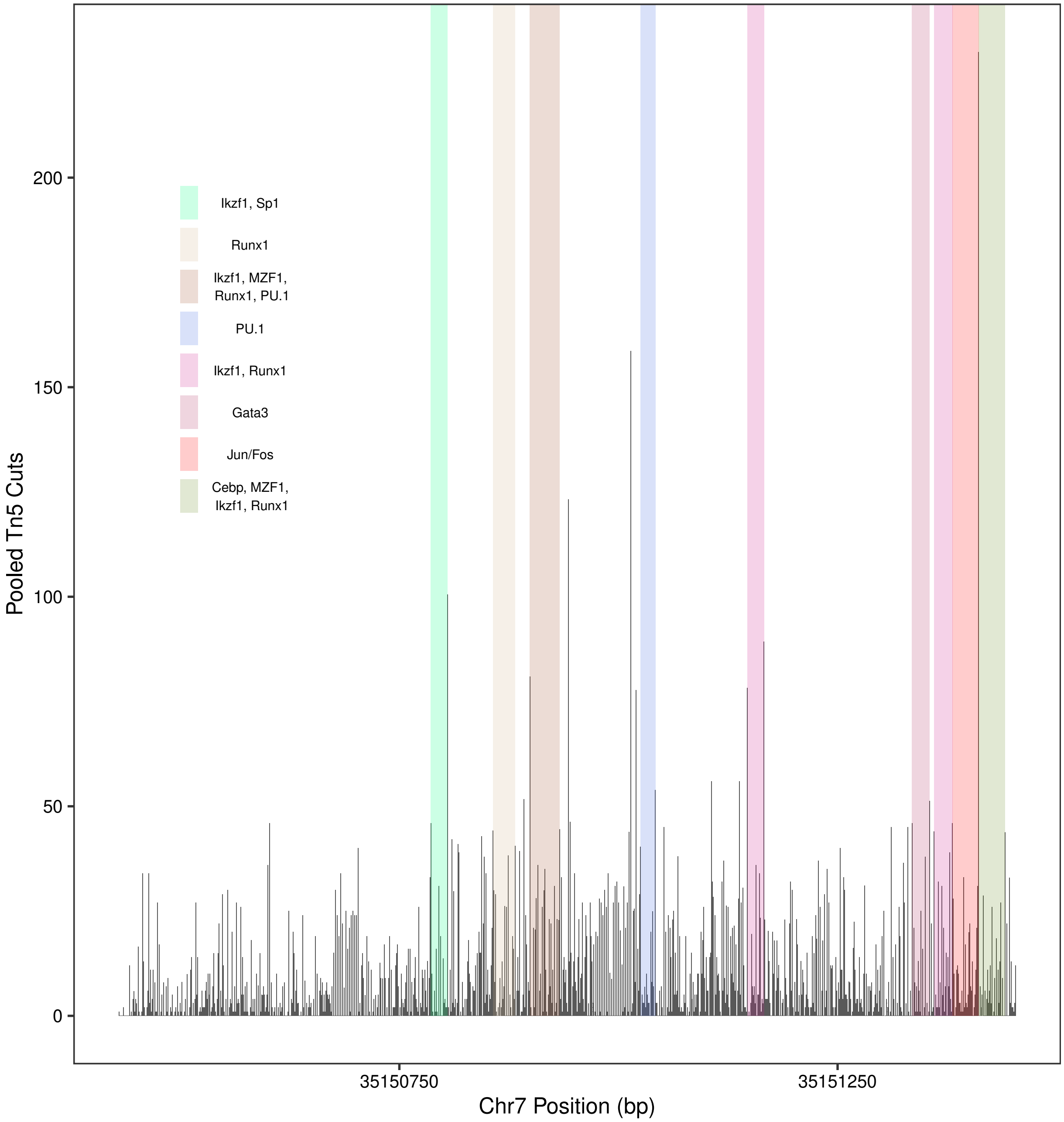
Ikzf1, Sp1
Runx1
Ikzf1, MZF1, Runx1, PU.1
PU.1
Ikzf1, Runx1
Gata3
Jun/Fos
Cebp, MZF1, Ikzf1, Runx1
Pooled Tn5 Cuts
0
50
100
150
200
35150750
35151250
Chr7 Position (bp)

Figure S9

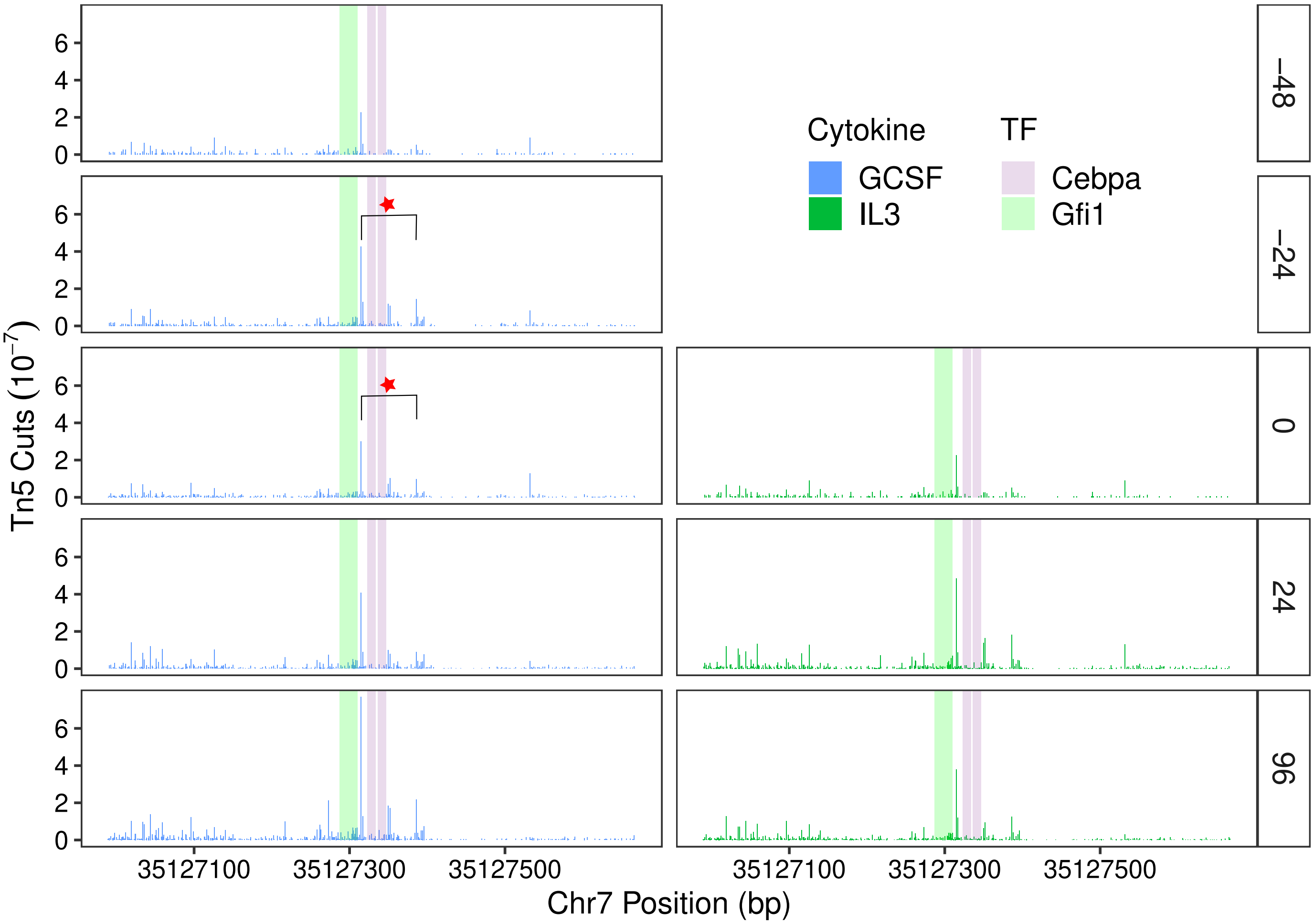

Tn5 Cuts ($10^{-7}$)
Chr7 Position (bp)
Cytokine
GCSF
IL3
TF
Cebpa
Gfi1
−48
−24
0
24
96
35127100
35127300
35127500


Figure S10

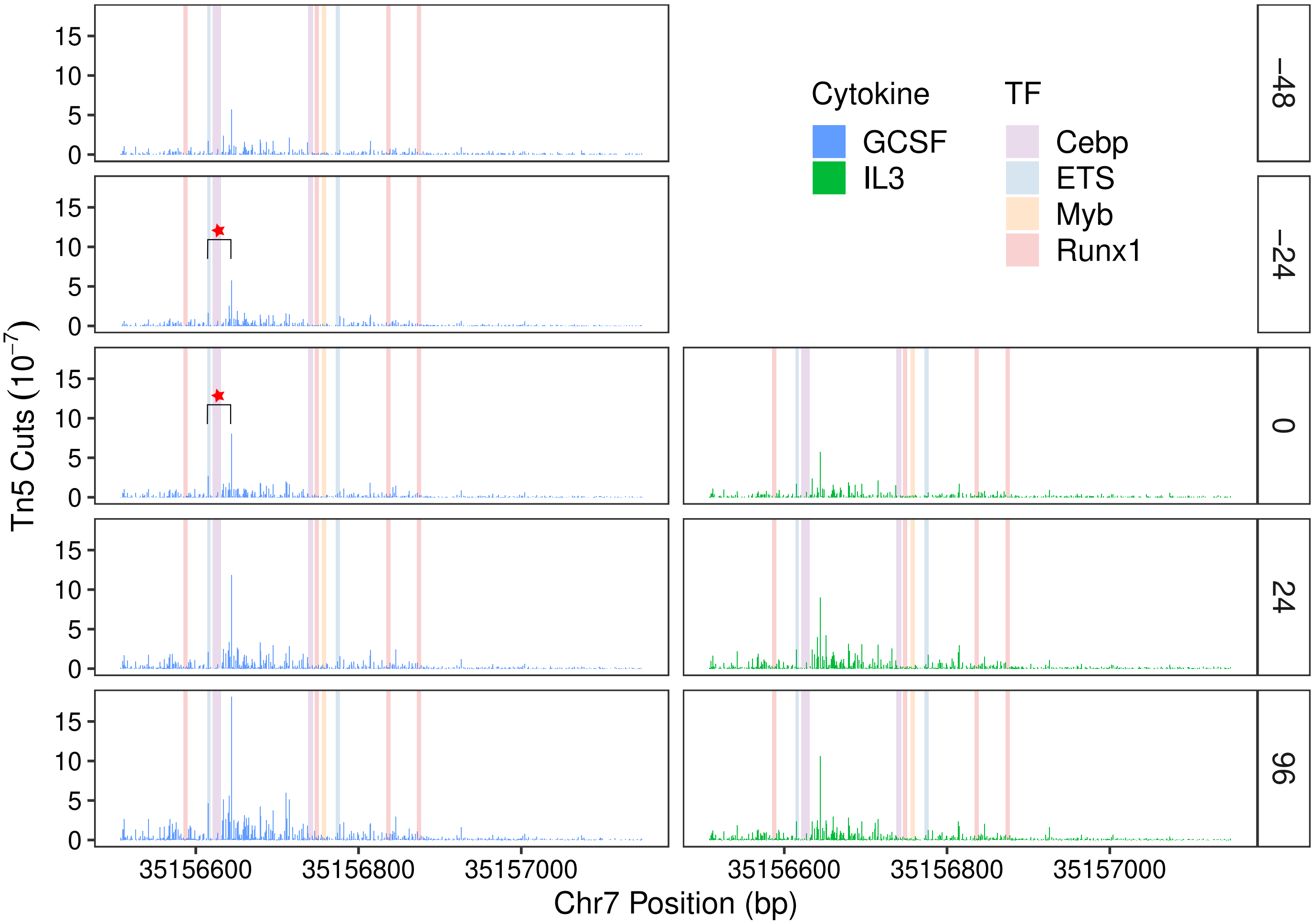
Cytokine
GCSF
IL3
TF
Cebp
ETS
Myb
Runx1
−48
−24
0
24
96
Tn5 Cuts ($10^{-7}$)
Chr7 Position (bp)
35156600
35156800
35157000
0
5
10
15

Figure S11

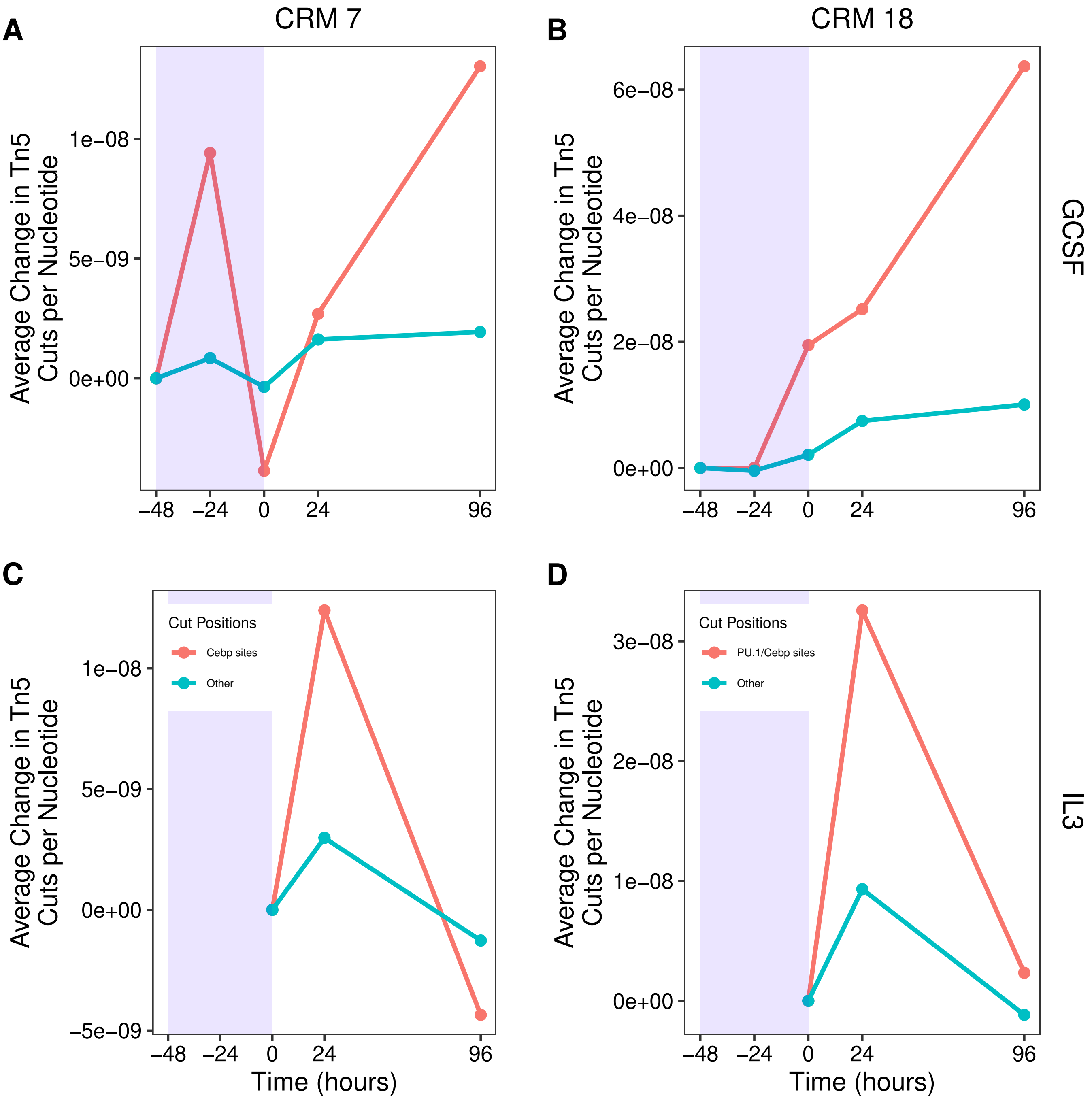
A
CRM 7
B
CRM 18
C
D
GCSF
IL3
Average Change in Tn5 Cuts per Nucleotide
Time (hours)
Cut Positions
Cebp sites
PU.1/Cebp sites
Other
1e−08
5e−09
0e+00
−5e−09
6e−08
4e−08
2e−08
3e−08
−48
−24
0
24
96

Figure S12

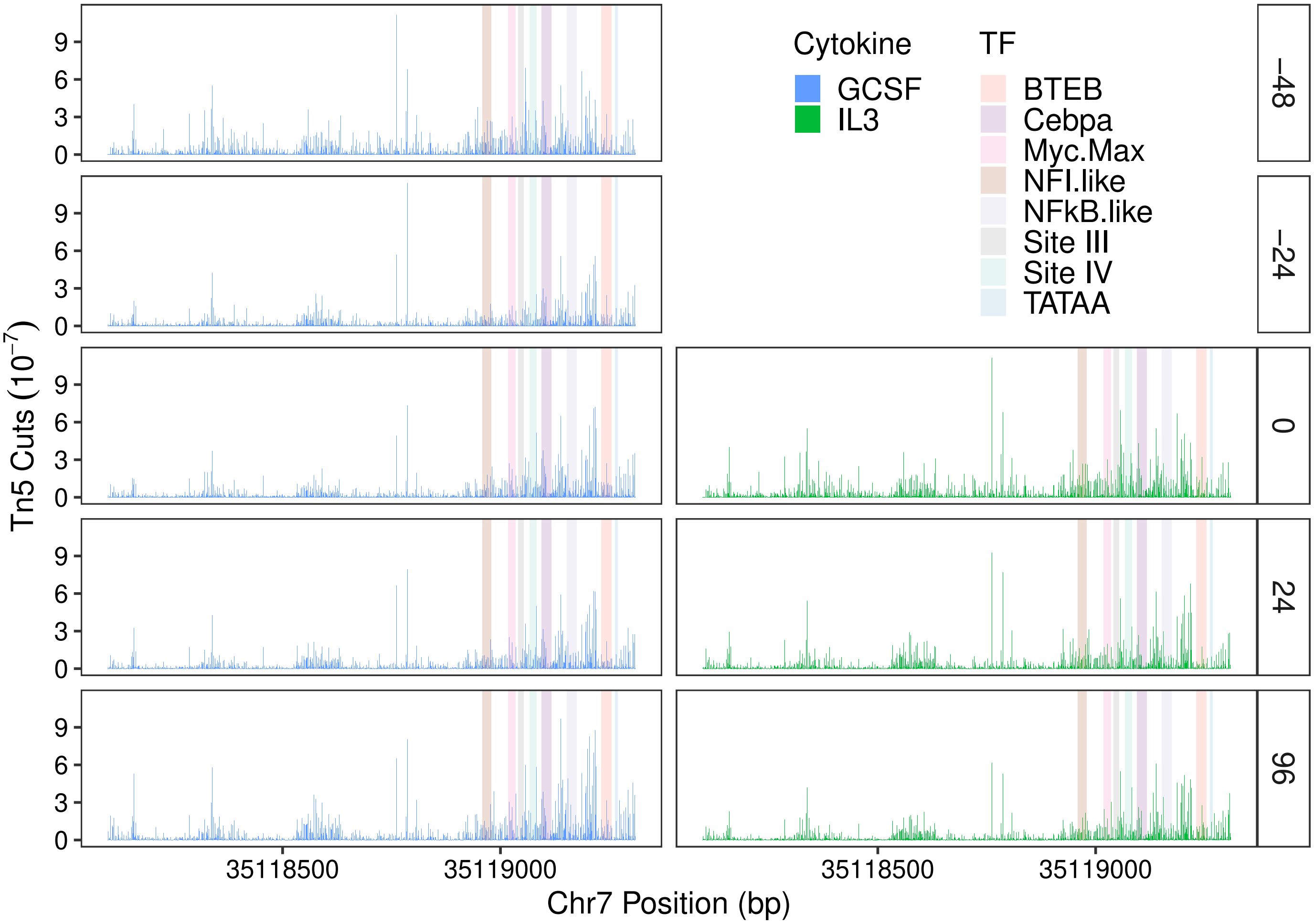
Cytokine
GCSF
IL3
TF
BTEB
Cebpa
Myc.Max
NFI.like
NFkB.like
Site III
Site IV
TATAA
−48
−24
0
24
96
Tn5 Cuts ($10^{-7}$)
35118500
35119000
Chr7 Position (bp)

Figure S13

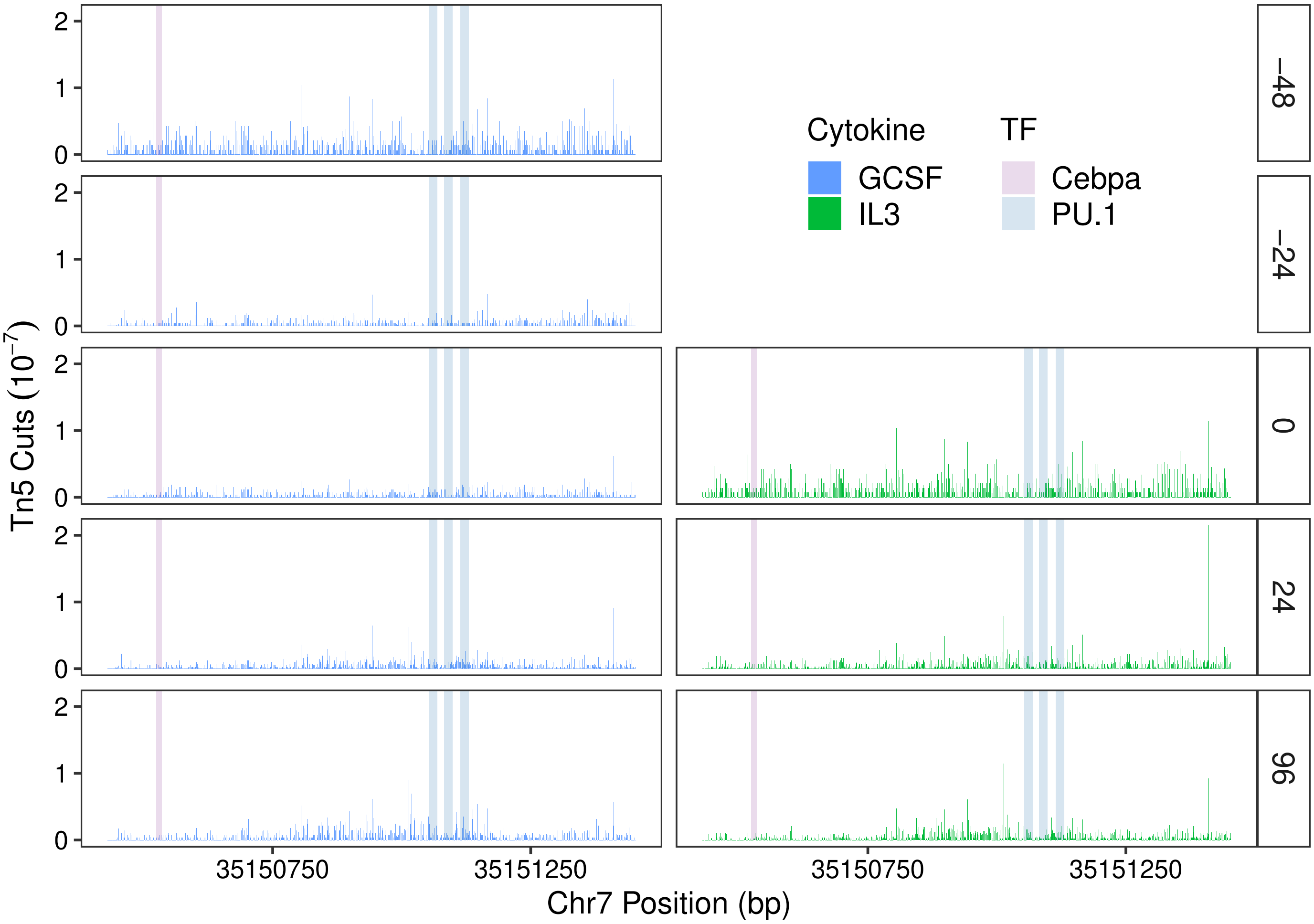
Tn5 Cuts (10^−7)
Cytokine
GCSF
IL3
TF
Cebpa
PU.1
−48
−24
0
24
96
0
1
2
35150750
35151250
Chr7 Position (bp)

Figure S14

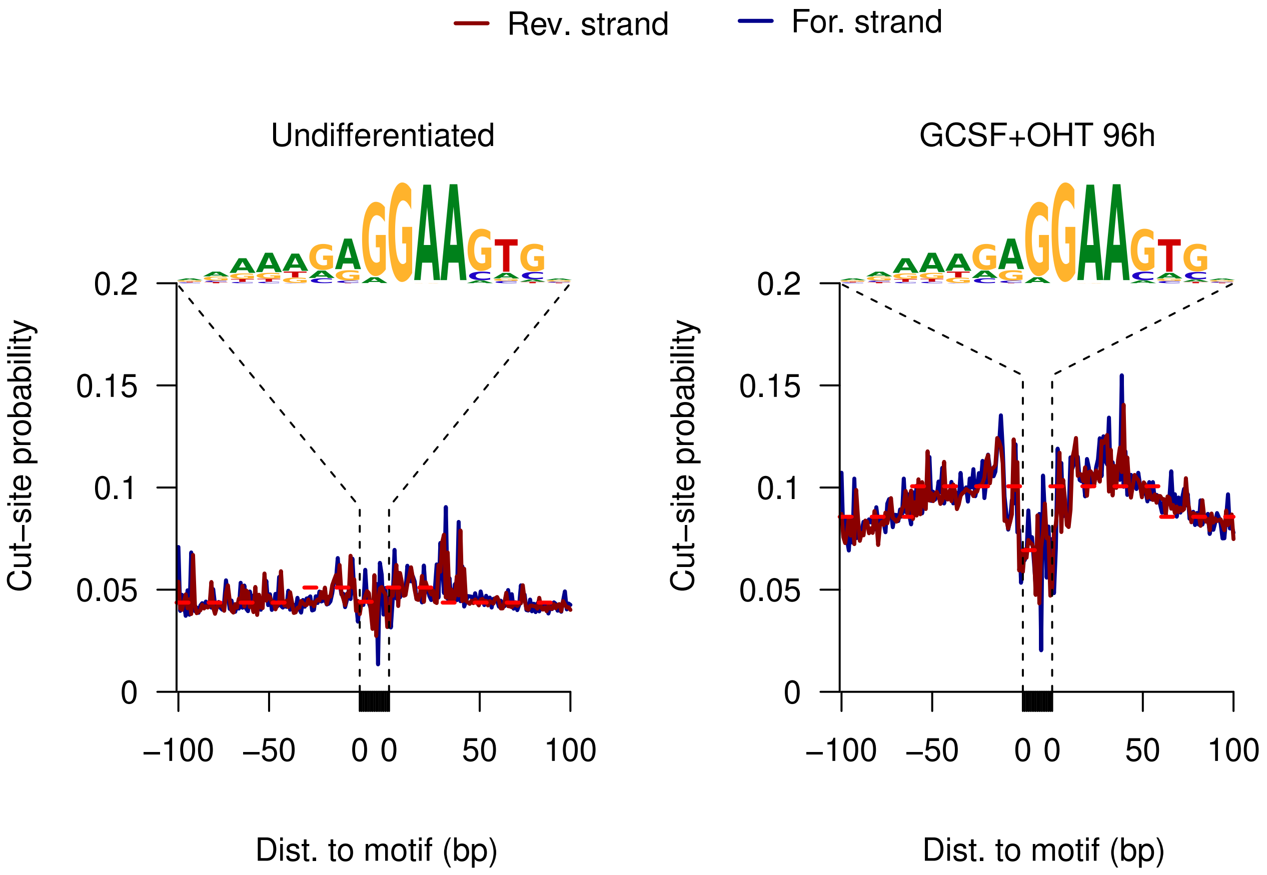
Rev. strand
For. strand
Undifferentiated
GCSF+OHT 96h
Cut–site probability
0.2
0.15
0.1
0.05
0
–100
–50
0 0
50
100
Dist. to motif (bp)

Figure S15

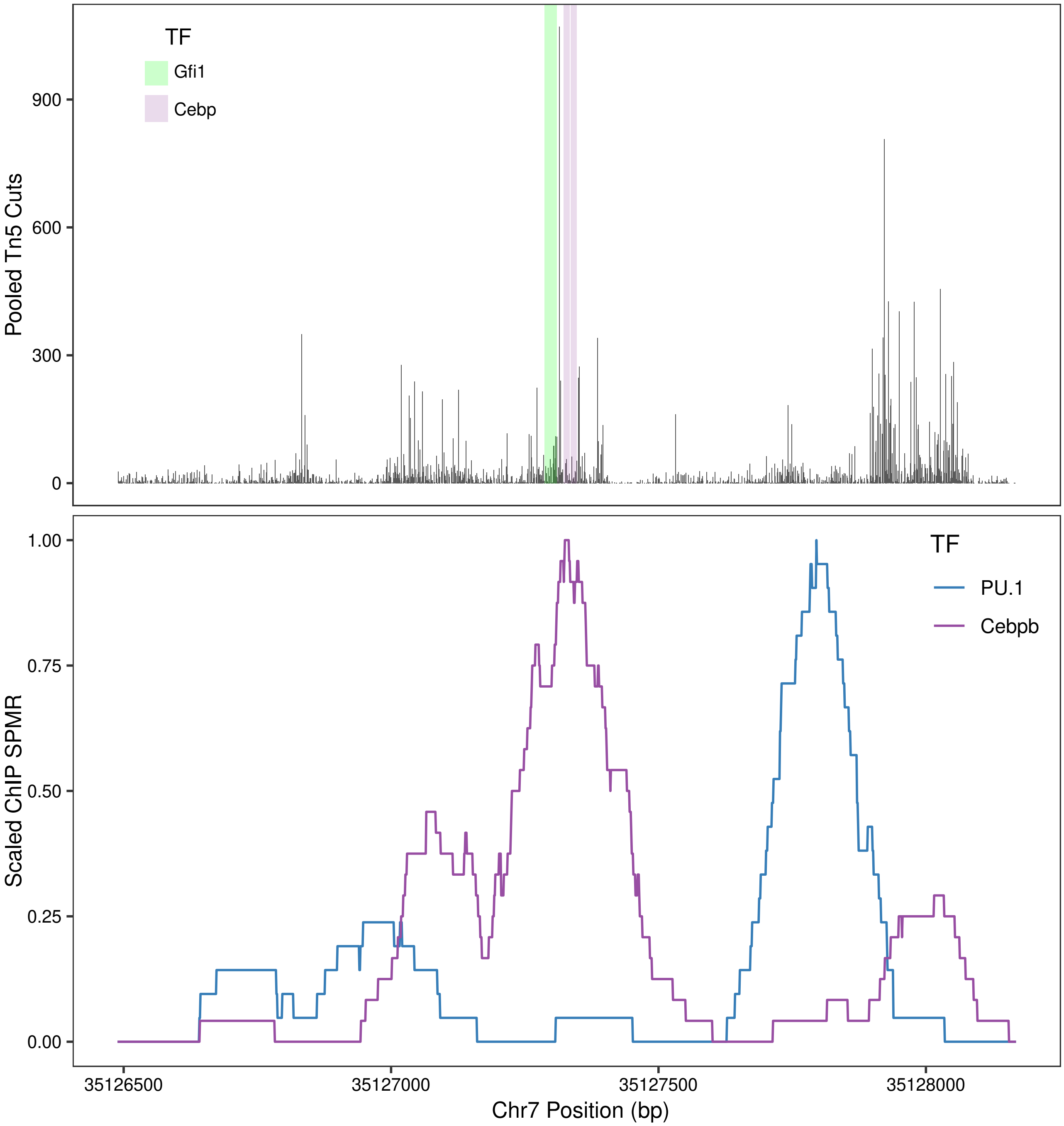
TF
Gfi1
Cebp
900
600
300
0
Pooled Tn5 Cuts
1.00
0.75
0.50
0.25
0.00
Scaled ChIP SPMR
TF
PU.1
Cebpb
35126500
35127000
35127500
35128000
Chr7 Position (bp)

Figure S16
